# Chemically Induced Degradation of Native Proteins by Direct Recruitment to the 26S Proteasome

**DOI:** 10.1101/2023.07.19.549534

**Authors:** Madeline Balzarini, Weijun Gui, Isuru M. Jayalath, Bin-Bin Schell, Joel Tong, Thomas Kodadek

## Abstract

Targeted protein degradation (TPD) is a promising strategy for drug development. Most degraders function by forcing the association of the target protein (TP) with an E3 Ubiquitin ligase, which in favorable cases results in the poly-Ubiquitylation of the TP and its subsequent degradation by the 26S proteasome. Here we explore the feasibility of a different TPD strategy in which the TP is recruited directly to the proteasome without the requirement for poly-Ubiquitylation. Using an engineered cell line in which the HaloTag protein is fused to one of the Ubiquitin receptors, we show that native protein targets can be degraded in this fashion when the cells are exposed to a chemical dimerizer containing a chloroalkane and a TP ligand. The potential advantages and disadvantages of Ubiquitin-independent degraders vs. traditional proteolysis-targeting chimeras are discussed.

---

Targeted protein degradation (TPD) has emerged as an attractive strategy for the development of drug candidates and tool compounds. (1) TPD employs chemical dimerizers known as PROTACs (2–4) (proteolysis-targeting chimeras) or molecular glues (5) that mediate the association of a target protein (TP) with an E3 Ubiquitin (Ub) ligase. In favorable cases, this results in the poly-Ubiquitylation of the TP and its subsequent degradation by the 26S proteasome.

A major advantage of PROTACs and molecular glues is that they function via an event-driven mechanism of action (MoA). In other words, degraders act catalytically. (6) This greatly reduces the bar for the type of chemical matter one requires to achieve potent inhibition of the TP relative to traditional inhibitors that function by an occupancy-driven MoA, because achieving a long residence time of the inhibitor on the TP is not necessary. However, current TPD modalities have limitations. It is clear that forced proximity of the E3 Ub ligase and the TP is necessary, but not sufficient, for efficient TP degradation. (7,8) Significant neo-protein-protein interactions (neo-PPIs) are necessary to stabilize the ternary complex, and a suitable alignment of an acceptor lysine residue on the TP and the activated Ub molecule must be achieved. (9) This results in the requirement for extensive linker optimization for the development of potent PROTACs. (10) Perhaps more problematic for oncology application is the prediction that resistance to PROTACs is likely to arise rapidly through mutation or down-regulation of the E3 Ub ligase, most of which are not essential proteins. (11,12)

Given these issues, and the prevailing view that poly-Ubiquitylation mainly serves as a signal to traffic the TP to the proteasome, we became interested in the development of Ub-independent degraders (UIDs) that function by delivering the TP directly to the 26S proteasome. Most of the subunits of the proteasome are encoded by essential genes, so it may be difficult for cancer cells to generate resistance to a Ub-independent degrader by simply reducing the expression of a proteasome component. In addition, there would be no requirement for a specific geometric orientation of the TP with an activated Ub molecule, so it is possible that linker optimization in a Ub-independent PROTAC would be straightforward.

There is ample precedent suggesting that the development of UIDs is possible. Several proteasome substrates, such as ornithine decarboxylase (ODC), are degraded by the proteasome without the requirement for poly-Ubiquitylation, thanks to the presence of proteasome-binding motifs in the protein that, when exposed, allow the substrate to be trafficked to the proteasome. (13,14) Moreover, several studies have shown that even proteins normally degraded via a Ubiquitin-dependent pathway can bypass the requirement for this post-translational modification if they are delivered to the proteasome by some other means. For example, pioneering work in yeast by the Church and Finley laboratories demonstrated that by fusing the FPR1 protein to a component of the 26S proteasome and the Tor protein to a model substrate, proteasome-dependent substrate turnover of the target protein was observed when rapamycin, which heterodimerizes Tor and FPR1, was added to the cells. (15) Matouschek and co-workers reported a similar result in mammalian cells (16) and furthermore showed that genetic fusion of target proteins to proteasome-associated proteins dramatically reduced their half-lives, (17) presumably by trafficking them to the proteasome. All these experiments strongly support the idea that proteasome localization, not poly-Ubiquitylation per se, is sufficient for protein turnover so long as the substrate has an unstructured region that allows it to be engaged by the AAA ATPases that unwind the substrate and feed it into the catalytic pocket of the 20S core particle. (18)

The feasibility of creating UIDs was recently demonstrated emphatically by a group at Genentech. (19) They discovered a large, macrocyclic peptide ligand for recombinant Rpn1 (PSMD2), a component of the proteasome’s 19S regulatory particle (19S RP) that recognizes a pocket not occluded by association of the protein with the proteasome. A conjugate of this macrocycle to a high affinity BRD4 ligand was shown to be slightly cell permeable and able to trigger the proteasome-mediated degradation of BRD4.

These studies set the stage for development of drug-like degraders. A central challenge is that few, if any, drug-like small molecules exist that bind the 19S Regulatory Particle (19S RP) of the proteasome, but are not proteasome inhibitors. (20–23) Therefore, a significant ligand discovery effort will be required to advance this field. An important decision in this effort will be what 19S RP protein(s) to focus on as screening targets. Rpn13 is an attractive candidate for several reasons. First, Rpn13 is a ubiquitin receptor. (24) Thus, using Rpn13 as a point of recruitment for ubiquitin-independent directed proteolysis would be hijacking Rpn13 to do essentially the same thing it does naturally. Additionally, Rpn13 is non-essential in healthy cells (25) but appears to promote the proliferation of several types of cancer cells. (26–31) This may reduce the chance that perturbing Rpn13 by binding it or by blocking engagement of its native substrates would result in high toxicity in healthy cells while still preventing cancer cells from developing resistance by mutation or downregulation. Finally, Rpn13 is ligandable by small molecules, though existing ligands lack the appropriate characteristics to be useful in the construction of UIDs. (20,23,32) However, a potential drawback of targeting Rpn13 for UID development is that the protein is not proximal to the AAA ATPases in the 19S RP complex, (33) possibly necessitating the use of long linkers to join Rpn13 and target protein ligands.

In this study we probe the suitability of Rpn13 as a target for UIDs. Specifically, we describe the creation of a cell line in which the proteasome-binding Pru domain of Rpn13 is fused to HaloTag protein (HTP). (34) We hypothesized that chimeric molecules containing a chloroalkane and a target protein ligand will act as UIDs in this cell line. Indeed, we show that such chimeras containing JQ1, (35) a bromodomain ligand, trigger the proteasome-mediated degradation of BRD2 protein without the need for Ubiquitylation. As anticipated, we show that the potency of these UIDs is linker length-dependent, with longer linkers providing more efficient BRD2 degradation. However, in contrast to traditional PROTACs, there is no sharp optimum for linker length. These studies validate Rpn13 as a potential target for UIDs.

## Results

### Establishment of a stable cell line enabling UID-mediated degradation of native proteins

Suitable ligands for Rpn13 that would allow it to be evaluated as a point of recruitment for ubiquitin independent degradation do not exist. To overcome this limitation, HaloTag7 protein (HTP) fusions to intact Rpn13 or domains of the protein were created. HTP reacts covalently with primary chloroalkanes, (34) which thus can serve as a surrogate ligand of endogenous Rpn13. Rpn13 has three major regions; the PRU domain near the N-terminus, which docks with Rpn2 in the 19S RP and recognizes ubiquitin, an unstructured central region, and the DEUBAD domain at the C-terminus, which binds and activates UCH37. (36–39) Since the PRU domain is closest to the AAA ATPases and is essential for proteasome binding, we decided to focus on HTP-PRU fusion constructs. Specifically, an expression vector was created encoding Rpn13 residues 1-128 followed by a C-terminal HTP and FLAG tag. Rpn13(1–128)-HTP-FLAG was then stably overexpressed in HEK293 cells using the Flp-In system by Invitrogen. In addition, a cell line stably expressing HTP-FLAG not fused to the Rpn13 PRU domain was created as a control. **Figure 1A** shows a schematic representation of the overexpressed proteins.

**Figure 1:**
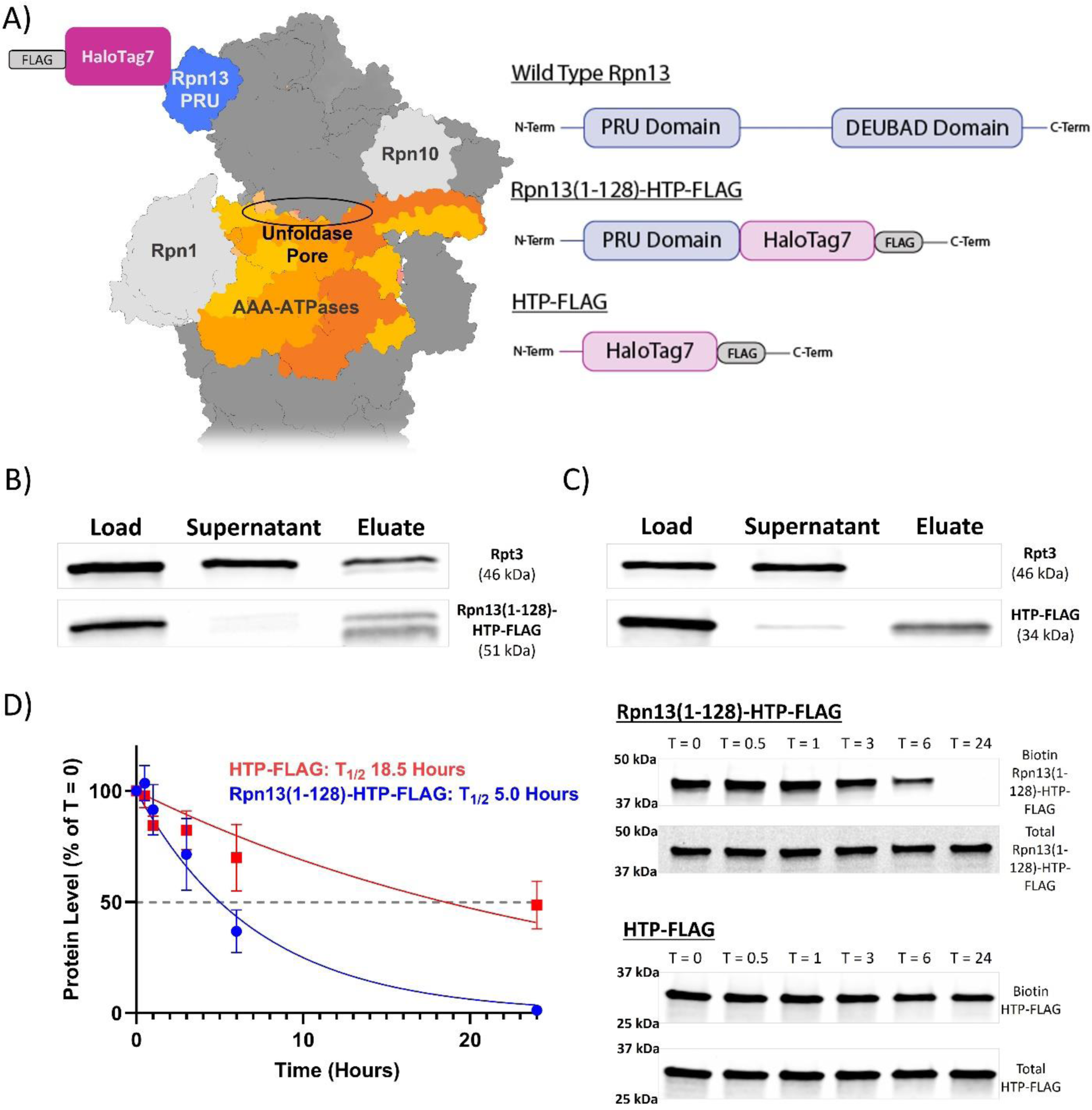
Characterization of Rpn13(1-128)-HTP-FLAG. A) Schematic view of the placement of the HaloTag7 protein, which is fused to the PRU domain of Rpn13. fusion was added to the Rpn13 PRU domain to act as a point of recruitment for proteasomal degradation. B) Analysis of the association of Rpn13(1–128)-HTP-FLAG with the 19S RP as determined by co-immunoprecipitation. Rpn13(1–128)-HTP-FLAG was immunoprecipitated from a cell lysate using immobilized anti-FLAG antibody. The presence of Rpt3 in the precipitate was evaluated by SDS-PAGE and Western blotting using anti-Rpt3 antibody. C) Analysis of the association of HTP-FLAG with the 19S RP using the same protocol described in B). D) Half-life of HaloTag7 fusion proteins as measured by a biotin-chloroalkane pulse/chase assay (see text for details). Right: Representative SDS-PAGE analysis of the level Rpn13(1–128)-HTP-FLAG or HTP-FLAG at various times after washout of excess biotin-chloroalkane. Left: Graph of the gel data shown on the right. The error bars represent the results from three independent experiments.

To determine if Rpn13(1–128)-HTP-FLAG associates with the proteasome, a lysate from cells expressing this construct, as well as a lysate from cells expressing the HTP-FLAG control protein, were prepared, and incubated with immobilized anti-FLAG antibody. The immunoprecipitants were analyzed by Western blotting using an anti-Rpt3 (one of the proteasomal ATPases) antibody (**Figure 1B-C**). Rpt3 co-precipitated with Rpn13(1–128)-HTP-FLAG and did not co-precipitate with HTP-FLAG. This demonstrates Rpn13(1–128)-HTP-FLAG is associated with the 19S RP. As expected, not all of the cellular Rpt3 was co-precipitated with Rpn13(1–128)-HTP-FLAG. These cells express endogenous Rpn13. Therefore, some percentage of the proteasomes contain the native Rpn13 and would not co-precipitate with Rpn13(1–128)-HTP-FLAG.

A concern with this experimental design is that proteasome-associated Rpn13(1–128)-HTP-FLAG might be degraded rapidly if an unstructured region can be engaged by the AAA ATPases. Since PROTAC-mediated degradation typically takes place over several hours, this is an important issue. Thus, the half-life of the protein was measured using a pulse-chase protocol. Cells expressing Rpn13(1–128)-HTP-FLAG or HTP-FLAG were treated with a chloroalkane-biotin conjugate (biotin-ct) for 1 hour to alkylate the HTP. Unreacted biotin-ct was then washed out and chased with chloroalkane tagged tryptophan (tryptophan-ct) which is highly cell permeable. At various times after biotin-ct washout, cell lysates were prepared and analyzed for biotinylated protein by Western blot. As shown in **Figure 1D**, the half-life of Rpn13(1–128)-HTP-FLAG was 5 hours, while that of HTP-FLAG was 18.5 hours. These data show that, as anticipated, localization of the HTP to the proteasome via the PRU domain fusion indeed stimulates its turnover, but we reasoned that this five-hour half-life was sufficient to support our experiments.

### Degradation of BRD2

With a suitable cell line in hand, a family of UID candidates targeting the bromo and extra-terminal domain (BET) family proteins (40) was synthesized by linking the bromodomain ligand JQ1 (35) to a chloroalkane via linkers comprised of one to six ethylene glycol units (**Figure 2A**; see SI for details of the synthesis and characterization). Conditions were then established for the efficient alkylation of Rpn13(1–128)-HTP-FLAG by these molecules *in cellulo*. Cells expressing Rpn13(1–128)-HTP-FLAG were treated with a JQ1-PEG-chloroalkane conjugate (10 µM), henceforth called a Halo-UID, for 30 minutes. The cells were washed and immediately lysed in the presence of a biotin-chloroalkane conjugate, which reacts with any Rpn13(1–128)-HTP-FLAG that was not alkylated by the Halo-UID molecule during the 30-minute incubation. The lysates were then analyzed by blotting with labeled Streptavidin (SA) to determine the amount of Rpn13(1–128)-HTP-FLAG biotinylation. These data (**Figure 2B**) showed that essentially 100% of the Rpn13(1–128)-HTP-FLAG inside the cell reacts with the Halo-UID during the 30-minute incubation, regardless of linker length.

**Figure 2:**
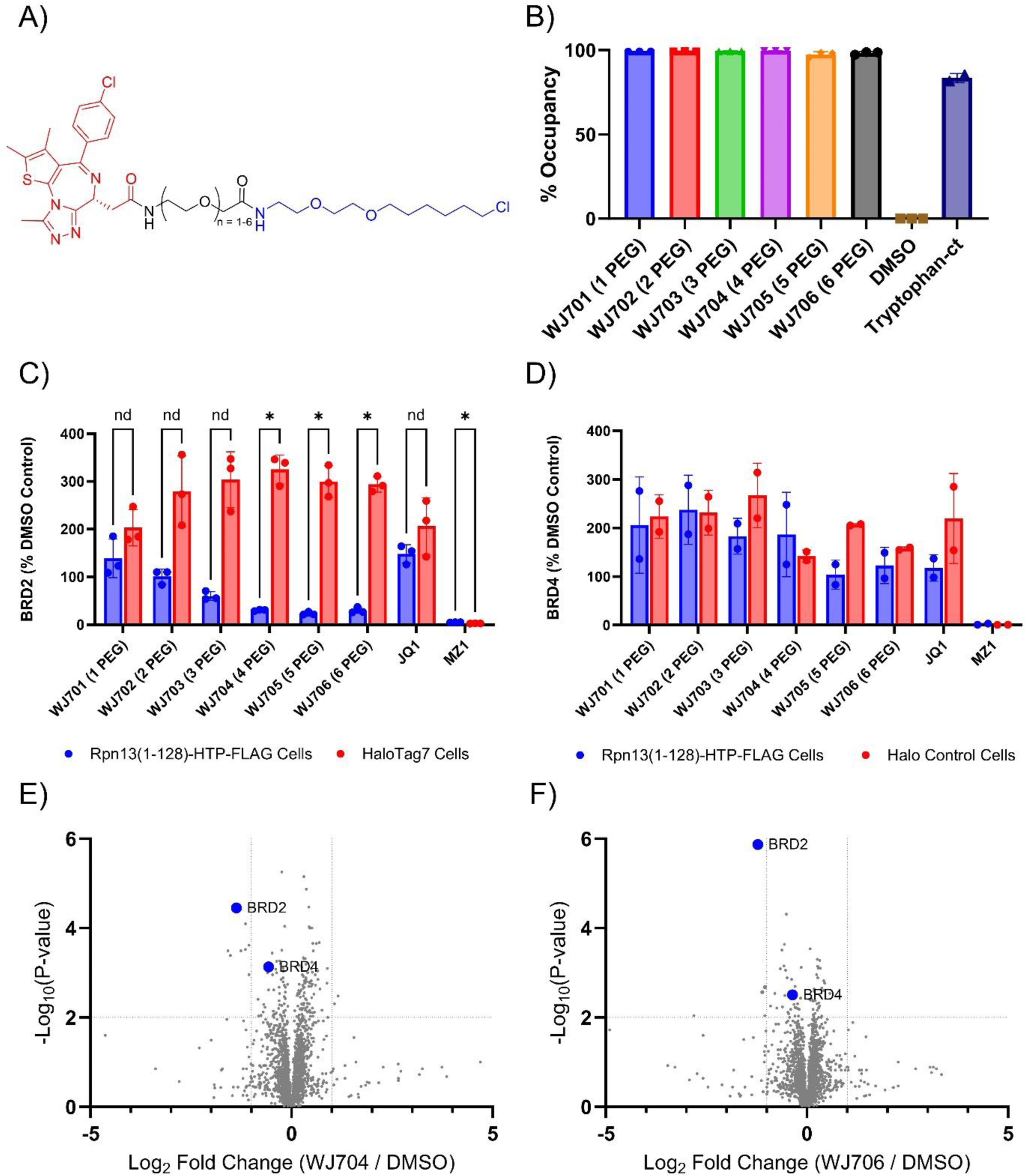
Selective Degradation of BRD2. A) The general structure of the BET targeting Halo-UIDs, with linkers range from 1-6 ethylene glycol units. B) Evaluation of the degree of Rpn13(1–128)-HTP-FLAG alkylation by the Halo-UIDs after a 30-minute incubation, as determined by subsequent labeling with a chloroalkane-biotin conjugate and analysis of the degree of Rpn13(1–128)-HTP-FLAG biotinylation by SDS-PAGE and blotting with labeled Streptavidin. The data are plotted as the percentage of the protein that was protected from biotinylation in three biological replicates. C) Assessement of BRD2 levels in cells that express Rpn13(1–128)-HTP-FLAG (blue) or HTP-FLAG (red) four hours after treatment with the indicated Halo-UIDs, JQ1 (not linked to a chloroalkane), or the previously reported, ubiquitin-dependent BET protein degrader MZ1 (1 μM, no washout). Protein levels were determined by SDS-PAGE and Western blotting. D) Assessment of BRD2 levels in cells that express Rpn13(1–128)-HTP-FLAG (blue) or HTP-FLAG (red) four hours after treatment with the indicated Halo-UIDs, JQ1 (not linked to a chloroalkane), or the previously reported, ubiquitin-dependent BET protein degrader MZ1 (1 μM, no washout). Protein levels were determined by SDS-PAGE and Western blotting. E. Global proteomics analysis of Rpn13(1–128)-HTP-FLAG-expressing cells treated with the indicated compounds. See text for details.

The ability of the Halo-UIDs to degrade endogenous BETs was assessed in the HTP expressing cell lines. Cells were treated with the Halo-UID (10 µM) for 30 minutes. Unreacted compound was then washed away. This protocol was designed to prevent excess, unreacted JQ1-chloroalkane conjugate from competing with proteasome-displayed JQ1 for binding to the BRD proteins. The cells were incubated for an additional four hours, after which lysates were prepared and the levels of BRD2 and BRD4 were assessed by Western blotting (**Figure 2C-D**).

Treatment of cells with unmodified JQ1 (10 μM) using this protocol resulted in increased BRD2 levels as compared to the DMSO control in both Rpn13(1–128)-HTP-FLAG and HTP-FLAG expressing cells, reflecting a known compensatory mechanism by which the cell adapts to BRD2 protein inhibition by producing more protein. (41) In contrast, incubation of this cell line with Halo-UIDs WJ702 – WJ706 (containing 2-6 ethylene glycol units in the linker) resulted in significantly decreased levels of BRD2. This was not the case in cells incubated with WJ701, which contains the shortest linker. Indeed, the activity of the Halo-UIDs increased with linker length up to four ethylene glycol units, after which a further increase did not increase or decrease the activity of the degrader. None of the Halo-UIDs decreased BRD2 levels in cells expressing HTP not linked to Rpn13. These data are consistent with the idea that Halo-UID-mediated trafficking of BRD2 to the 19S RP results in its degradation so long as the linker length is sufficient to allow the AAA ATPases to “reach” the substrate.

JQ1 is known to engage other BET proteins including BRD4. (35) Therefore, the effect of the Halo-UIDs on BRD4 levels was also assessed.

BRD4 levels were not decreased by treatment with any of the Halo-UIDs (**Figure 2D**). In all cases, an increase in protein levels relative to the DMSO control was observed, as was also the case for JQ1 treatment. This again represents the cell’s response to BRD4 inhibition.

Proteomics was used to verify the selectivity of the BET targeting Halo-UIDs. Two of the BRD2 degrading compounds, WJ704 (4 PEG linker) and WJ706 (6 PEG linker), were selected for analysis. Samples were run in quadruplicate and analyzed by TMT-based global proteomics. (42) Unique peptides for BRD2 and BRD4 were detected. ANOVA analysis with a false discovery rate of 1% and a significance level of p = 0.01 with multiple hypothesis corrections indicated the levels of only two proteins were significantly changed upon compound treatment, BRD2 and WD repeat containing protein 75 (WDR75). Pairwise comparisons, shown in **Figure 2E-F**, indicate that BRD2 is the only protein significantly decreased by more than 1 log-fold with a p < 0.01. These data confirm the selective degradation of BRD2 by the Halo-UIDs.

As BRD4 has been successfully targeted for proteasomal degradation by several JQ1 based degraders, (41,43,44) the failure of the Halo-UIDs to induce BRD4 degradation was surprising. To probe this issue further, we asked if the Halo-UIDs indeed recruited BRD4 to the proteasome by carrying out co-immunoprecipitation experiments. Rpn13(1–128)-HTP-FLAG cells were pre-treated with bortezomib (10 μM) to block proteasomal degradation and then treated with Halo-UID (10 μM) for 30 minutes. Compounds were then washed out and the cells were incubated for two hours to allow time for complexes to form. Cells were lysed and Rpn13(1–128)-HTP-FLAG was precipitated using anti-FLAG antibody displaying resin. Western blot analysis of the input and precipitate, shown in **Figure 3**, indicate that both BRD2 and BRD4 co-precipitated with Rpn13(1–128)-HTP-FLAG in the presence of Halo-UIDs but not in the presence of the vehicle control. Co-precipitation of BRD2 with WJ701 (a non-degrading Halo-UID) and of BRD4 with all tested Halo-UIDs indicates preferential formation of the BRD2/Rpn13(1–128)-HTP-FLAG complex was not the cause of the BRD2 selective degradation. These data demonstrate that UID-mediated localization of a TP to the proteasome, while necessary, is not sufficient for efficient degradation.

**Figure 3:**
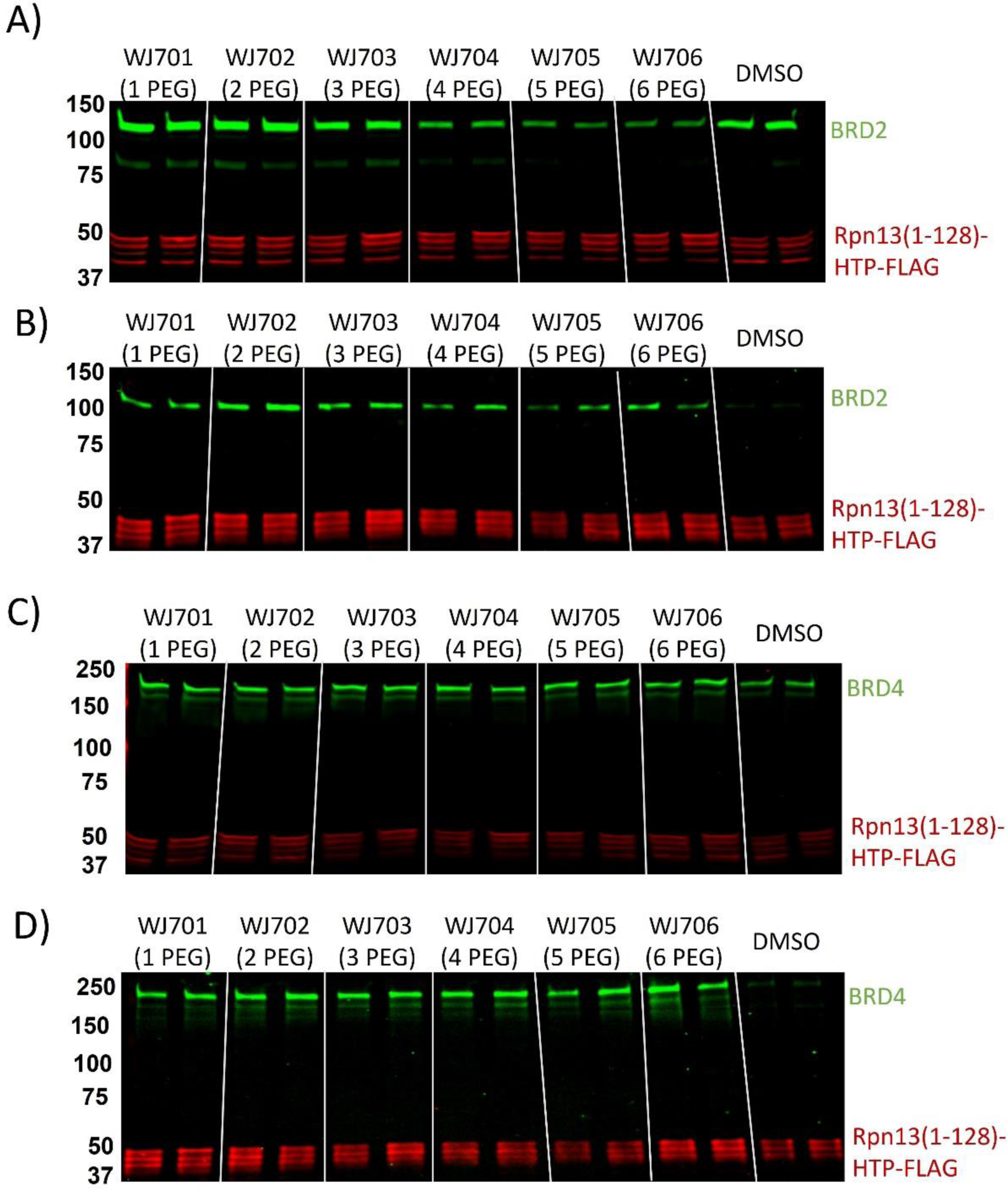
Halo-UID-mediated association of BRDs and Rpn13(1-128)-HTP-FLAG. Rpn13(1–128)-HTP-FLAG expressing cells were treated with Halo-UIDs (10 μM) and proteasome inhibitor. Cells were lysed and Rpn13(1–128)-HTP-FLAG was precipitated with anti-FLAG displaying resin; lysate and precipitate were analyzed by Western blotting for BRD2 or BRD4 and FLAG. A) BRD2 was clearly detected in the lysate after Halo-UID treatment, and B) BRD2 co-precipitated with Rpn13(1–128)-HTP-FLAG in the Halo-UID treated samples but not in the vehicle treated samples. Blotting for BRD4 and FLAG in the C) lysate and D) precipitate also shows co-precipitation of Rpn13(1–128)-HTP-FLAG and BRD4 upon Halo-UID treatment but not in vehicle treated cells. Two biological replicates for each sample are shown.

### BRD2 degradation does not require poly-ubiquitination

To determine if the observed decrease in BRD2 levels was the result of proteasomal degradation without prior ubiquitination, the effect of two ubiquitination inhibitors, TAK-924 and TAK-243, on the Halo-UID-mediated depletion of BRD2 was assessed.

TAK-924 is an inhibitor of NEDD8-activating Enzyme (NAE) and thus prevents the activation of cullin-RING E3 Ubiquitin ligases.(45) Halo-fusion expressing cells were co-treated with TAK-924 (10 μM) and WJ704, WJ705, or WJ706. Western blot analysis, shown in **Figure 4A-B**, revealed that NAE inhibition by TAK-924 did not have a significant impact on the degradation of BRD2 upon treatment with the Halo-UIDs. In stark contrast, BRD2 degradation by the VHL recruiting PROTAC, MZ1 (44) (1 μM) was blocked. These data confirm that the degradation of BRD2 by WJ704, WJ705, and WJ706 is independent of cullin-RING ligases.

**Figure 4:**
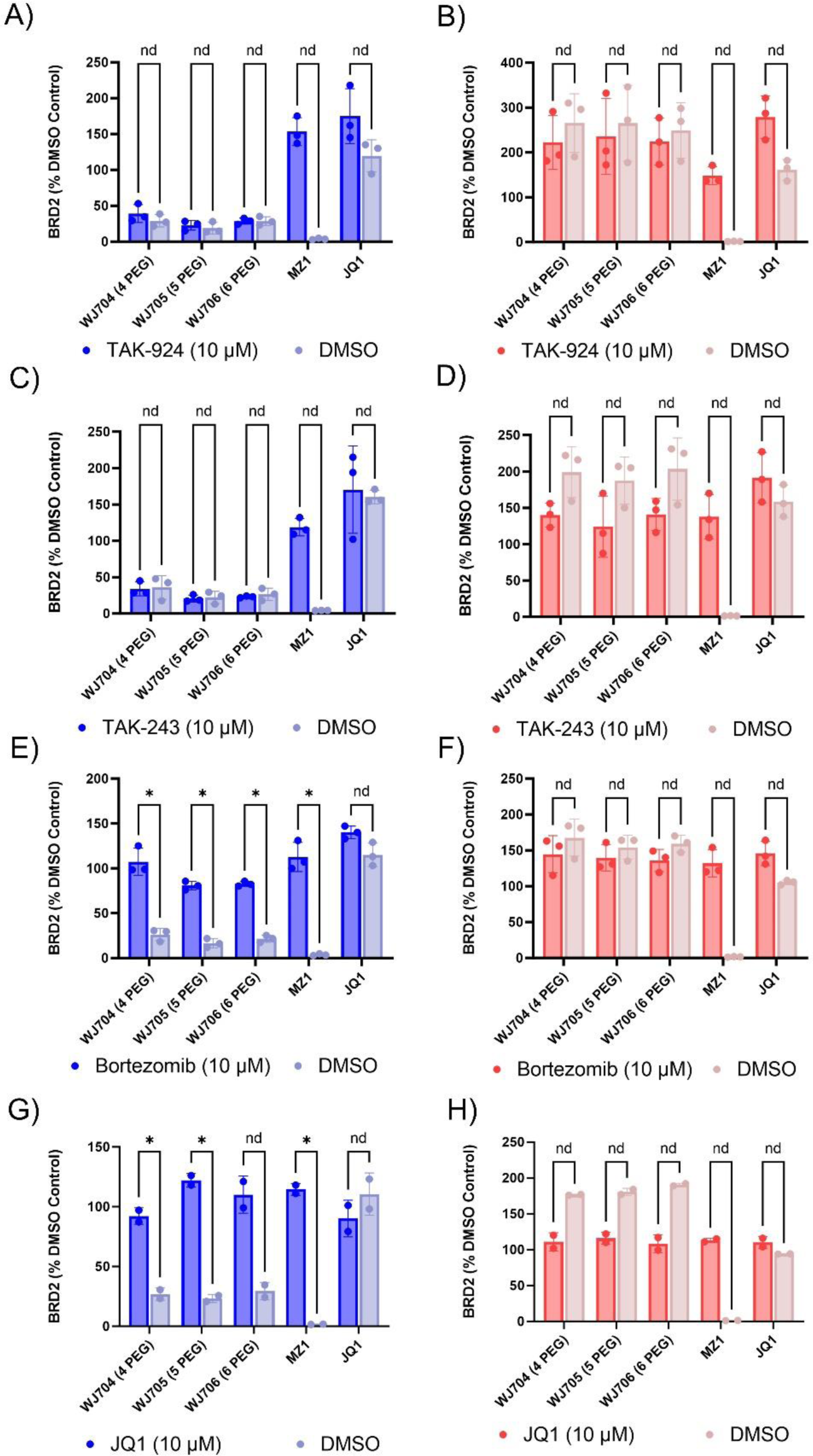
Halo-UID-mediated BRD2 degradation is insensitive to inhibitors of protein ubiquitylation but sensitive to proteasome inhibition. Cells expressing Rpn13(1–128)-HTP-FLAG (A, C, E and G; blue bar graphs) were treated with the indicated chemicals. The cells were lysed and the levels of BRD2 were assessed by SDS-PAGE and Western blotting. The bars represent the average of two - three biological replicates and the individual data points are shown as dots. A) Effect of the Nedd activating enzyme inhibitor TAK-924. C) Effect of the ubiquitin activating enzyme inhibitor TAK-243. E) Effect of the proteasome inhibitor Bortezomib. G) Effect of Added JQ1. The data show that the inhibitors of protein ubiquitylation had no effect on the activity of the three Halo-UIDs tested but blocked the activity of the ubiquitin-dependent degrader MZ1. Targeted degradation of BRD2 was blocked in all cases by the proteasome inhibitor Bortezomib or excess JQ1. The red panels (B, D, F and H) show the results of the same set of experiments in cells that express HTP-FLAG, which is not tethered to the proteasome.

To probe this issue in another way, the effect of TAK-243 (10 μM), a potent inhibitor of the Ubiquitin-activating enzyme UBA1, (46) on BRD2 degradation by the Halo-UIDs was examined. As shown in **Figure 4C-D**, blockade of Ubiquitin activation had no significant effect on the degradation of BRD2 by WJ704, WJ705, or WJ706 but blocked MZ1-mediated degradation completely.

To test the proteasome dependence of BRD2 degradation, the cells were treated with the inhibitor Bortezomib (10 μM) along with the Halo-UIDs. As shown in **Figure 4E-F**, Bortezomib blocked degradation of BRD2.

Finally, we verified that the recruitment of BRD2 to the proteasome for degradation was facilitated by specific binding to displayed JQ1 (**Figure 4G-H**). Cells were co-treated with JQ1 (10 µM) and Halo-UIDs for 30 minutes. Following compound washout, JQ1 (10 uM) was maintained in solution. As anticipated, Halo-UID induced BRD2 degradation in Rpn13(1–128)-HTP-FLAG cells was blocked by free JQ1 due to the free ligand competing for binding of BRD2 to proteasome-displayed JQ1. Degradation of BRD2 by MZ1 was also blocked by JQ1 competition in Rpn13(1–128)-HTP-FLAG and HTP-FLAG expressing cells.

Taken together, these data indicate that the activity of the Halo-UIDs is proteasome-dependent but not Ubiquitin-dependent, consistent with our proposed mechanism of degradation by direct recruitment to the 26S proteasome.

## DISCUSSION

We have demonstrated that endogenous BRD2 can be selectively degraded without prior ubiquitination when it is trafficked to the proteasome directly by a chemical dimerizer. Further, we have validated the ubiquitin receptor Rpn13 as a suitable association site for UIDs. These findings set the stage for the discovery of Rpn13-binding small molecules that do not compete with its association with the proteasome or inhibit any of its various activities.

The fact that BRD2, but not BRD4, was degraded by the UIDs described in this study was somewhat surprising. Our initial assumption was that, unlike traditional, Ubiquitin-dependent PROTACs, UID-induced proximity to the proteasome would be necessary and sufficient for TP degradation, so long as the linker connecting the chloroalkane and the TP ligand is long enough to allow an unstructured region of the TP to be engaged by the AAA ATPases. This is clearly not the case. The co-immunoprecipitation experiments shown in Fig. 3 reveal that both BRD2 and BRD4 are recruited to the proteasome by the UIDs described here, yet only BRD2 is degraded to a significant extent. We speculate that the orientation of BRD4 in the induced ternary complex may be such that its disordered region is not engaged by the ATPases, but understanding the nature of this selectivity will require further study.

## Experimental Methods

### Plasmid Cloning

HaloTag7 with a C-terminal FLAG Tag (HTP-FLAG) and Rpn13 amino acids 1-128 with c-terminal HaloTag7 fusion and FLAG tag (Rpn13(1–128)-HTP-FLAG) were cloned into the pcDNA5/FRT vector for use in Flp-In stable cell line generation. The Takara “In-Fusion” system was used. The vector backbone and the insert were amplified separately, assembled, and transformed into bacteria.

The pcDNA5-FRT backbone was amplified from pcDNA5-FRT-ADRM1-FLAG (addgene #19418). 10 ng of template DNA was combined with PrimeStarMax DNA Polymerase (Takara #R045Q) and 300nM primers and amplified using a BioRad T100 Thermal Cycler ([98°C, 10s; 52.7°C, 10s; 68*C, 120s] x 30 cycles). Amplified DNA was purified by preparative agarose gel using a NucleoSpin Gel and PCR Clean-up Kit (Machery-Nagel 740609.10).

The inserts were amplified from the HaloTag-PSMD3 fusion vector (Promega N2701). 10 ng of template DNA was combined with PrimeStarMax DNA Polymerase (Takara #R045Q) and 300nM primers and amplified using a BioRad T100 Thermal Cycler ([98°C, 10s; 61°C, 10s; 72°C, 20s] x 30 cycles). Amplified DNA was purified using a NucleoSpin Gel and PCR Clean-up Kit (Machery-Nagel 740609.10).

Assembly of the plasmid was completed according to manufacturer’s protocol. Briefly, In-Fusion Snap Assembly Master Mix (Takara #638951) was combined with PCR amplified insert and PCR amplified vector at a 2:1 molar ratio and heated at 50°C for 15 minutes. The assembled plasmid was carried forward without further purification.

Assembled plasmid was transformed into Stellar Cells (Takara #636766). 50 uL of competent cells were pre-incubated with 2.5 uL of the in-fusion reaction mix for 30 minutes on ice. Cells were heat-shocked 45 s at 42°C and then allowed to recover on ice for 2 minutes. Cells were grown out in 500 uL of SOC media at 37°C with shaking for 1 hour. The entire culture volume was then plated on an LB agar plate with 100 ug/mL ampicillin. Plates were grown overnight at 37°C.

Colonies were selected from the transformation plate and grown up overnight in 5 mL of LB + 100 ug/mL ampicillin. Plasmid DNA was isolated from the cultured bacteria using a QIAprep Spin Miniprep Kit (Qiagen # 27106). Plasmid DNA was sent to Genewiz for sequencing using the CMV-Forward and BGHR primers. Clones with the correct sequences were identified. Plasmid DNA was purified from 130 mL of liquid culture using a HiSpeed Plasmid Maxi Kit (Qiagen 12663).

### Flp-In 293 Cell Lines - Stable Selection

The newly constructed pcDNA5-FRT-HaloTag7-Flag and pcDNA5-FRT-Rpn13(1–128)-HTP-FLAG-FLAG plasmids were co-transfected with the pOG44 plasmid (Invitrogen #V6005-20) containing Flp recombinase into Flp-In 293 cells (Invitrogen R750-07) using Lipofectamine2000 (Thermo #11668019).

Flp-In 293 cells were plated in a 10 cm dish in complete media lacking antibiotics 24 hours in advance so they would be approximately 70% confluent at the time of transfection. pOG44 and either the pcDNA5-FRT-HaloTag7-Flag plasmid or the pcDNA5-FRT-Rpn13(1–128)-HTP-FLAG-Flag plasmid were combined at ratio of 9:1 (ng/ng) and transfected into Flp-In 293 cells according to the Lipofectamine2000 protocol.

24 hours after transfection the media was refreshed with complete media (DMEM + 4.5 g/L glucose, 2 mM L-Glutamine + 10% Heat Inactivated Fetal Bovine Serum) lacking antibiotics. 48 hours after transfection the cells were split 1:4 into four 10 cm dishes with complete media with 100 ug/mL Hygromycin (Thermo 10687010). The media was refreshed ever 3-4 days for the next 2 weeks while distinct colonies formed. Because the Flp-In 293 cells have a single site for FRT recombination, the colonies were assumed to be isogenic; all cells were collected and used to seed a T150 flask. The cells were grown to 90% confluence and used to seed multiple flasks. When they had reached 90% confluence the cells were collected for cryopreservation. Cells were pelleted and resuspended in complete media with 10% DMSO at a concentration of ∼1.5 x 10^7^ cells/mL. 1 mL aliquots were slow frozen and then stored in the liquid N_2_ vapor phase for later use.

### General Mammalian Cell Culture

Cells are propagated in complete media (DMEM + 4.5 g/L glucose, 2 mM L-Glutamine + 10% Heat Inactivated Fetal Bovine Serum) with 100 ug/mL hygromycin and incubate at 37°C in a humidified 5% CO_2_ atmosphere. Cells are sub-cultured 2-3 times per week; there are typically ∼307,000 cells/cm^2^ at confluence. When sub-culturing, the old media is aspirated off and the cells are washed gently with DPBS (no calcium, no magnesium). Cells are dissociated from the surface using TrypLE express (Thermo #12605010) and dilute with complete media to inactivate the trypsin. Cells can then be seeding into a fresh flask for propagation or into plates for experiments. Cells are grown out to passage 20.

### General Western Blotting Procedure

Cells were lysed in M-PER (Thermo 78501) + 1X Halt Protease Inhibitor (Thermo 1861279) according to the manufacturer’s protocol. Lysates were clarified by centrifugation at 14,000 RCF, 15 min, 4°C. Lysate concentration was measured by Coomassie Protein Assay (Thermo 1856209) and normalized to 2 mg/mL total protein. 23 ug of each sample in 1X laemmli buffer (BioRad 1610747) with β-mercaptoethanol were used for western blot analysis. 4-15% TGX gels were used when blotting for proteins that are more than 100 kDa, and 4-20% TGX gels were used when blotting for proteins less than 100 kDa. All gels were run at 180V in SDS-Tris-Glycine buffer, and Protein Kaleidoscope (BioRad 1610375) was used as a molecular weight marker. Protein was transferred to a 0.2 um Nitrocellulose membrane (BioRad 1704158) using a TransBlot Turbo semi-dry transfer system. Blots with target proteins less than 100 kDa were transferred using 1.3A, 25V, 7 min; blots with targets greater than 100 kDa were transferred using 1.3A, 25V, 15 min. Membranes were blocked for >60 min at room temperature with gentle shaking in 5% BLOT-QuickBlocker (w/v) (G-Bioscience 786-011) in phosphate buffered saline with 0.05% Tween-20 v/v (PBS-T).

The membranes were then probed overnight at 4°C with primary antibodies in 5% BLOT-QuickBlocker; a 1:1000 dilution of primary antibody in a total volume of 10 mL was used unless otherwise indicated. The blots were then washed 3x 15 min in PBS-T at room temperature and then probed with secondary antibodies or other detection reagents in Intercept blocking buffer (Licor 927-70001) for 1 hour at room temperature. Fluorescently labeled secondary antibodies for the detection of mouse (Licor 926-54012), rabbit (Licor 926-32211), and rat (Licor 926-68076) antibodies were used at a 1:10,000 dilution in a total volume of 10 mL unless otherwise indicated. After 3x 15 min washes in PBS-T, the membranes were imaged on a Licor Odyssey M scanner. Imagers were quantified using Empiria Studio.

### Proteasome Association

Incorporation of Rpn13(1–128)-HTP-FLAG into the 19S was assayed by co-immunoprecipitation. Rpn13(1–128)-HTP-FLAG and HTP-FLAG were precipitated using M2 anti-FLAG resin (Sigma #M8823) and the eluate was blotted for Rpt3, a member of the 19S.

Rpn13(1–128)-HTP-FLAG Cells and HTP-FLAG cells were lysed in M-PER (Thermo 78501) + 1X Halt Protease Inhibitor (Thermo 1861279) according to the manufacturer’s protocol. Lysates were clarified by centrifugation at 14,000 RCF, 15 min, 4°C. Lysate concentration was measured and normalized to 4 mg/mL with tris buffered saline (TBS). 10 uL of packed M2 resin was equilibrated into TBS. 400 ug of lysate was then added and tumbled with the resin for 2 hours at 4 °C. The resin was washed 3x with 500 uL of TBS, and the bound proteins were eluted by heating the resin in 1x Laemmli buffer at 95 *C for 10 minutes. Samples of the supernatant and the naïve lysate were also prepared for western blot analysis. Membranes were blotted for HaloTag7 (Promega G928A) and Rpt3 (Abcam 140515-1001). Co-precipitation of Rpt3 with Rpn13(1–128)-HTP-FLAG indicated that the fusion protein was incorporated into the 19S proteasome.

### Halo-Fusion Protein Half Life

The half-life of the stably expressed HTP-FLAG and Rpn13(1–128)-HTP-FLAG was measured using a pulse/chase assay. Cells were plated in a 6 well dish in complete media lacking antibiotics so that they would be 90% confluent at the beginning of the assay. Cells were treated with chloroalkane tagged biotin (50 nM) in complete media for 60 minutes to alkylate HaloTag fusion proteins. The compound containing media was removed, cells were washed with DPBS, and treated with chloroalkane tagged tryptophan (10 μM) in complete media to alkylate any newly synthesized HaloTag protein. The T = 0 timepoint was collected immediately, and subsequent timepoints were collected at 30 minutes, 1 hour, 3 hours, 6 hours, and 24 hours. Cell pellets were snap frozen at the time of collection. Western blots were run according to standard procedure and probed with streptavidin IRDye680RD (Licor 925-68079) to track biotinylated HaloTag protein and anti-HaloTag antibody (Promega G928A) to track total HaloTag protein. Streptavidin was added with the secondary antibody at a 1:1000 dilution in Intercept buffer supplemented with 0.2% Tween-20 and 0.1% Sodium Dodecyl Sulfate (SDS). Phosphate buffered saline with 0.2% Tween-20 and 0.1% SDS was also used as the wash buffer following streptavidin addition. The decrease in biotin signal was used to calculate the protein’s half-life.

### HaloTag Occupancy Assay

The fraction of Rpn13(1–128)-HTP-FLAG alkylated by putative BRD2 degraders was assessed. Cells were plated in a 6 well dish in complete media lacking antibiotics so that they would be 90% confluent at the beginning of the assay. Cells were treated with WJ701-706 (10 μM) in complete media lacking antibiotics for 30 minutes, during which time the compounds alkylated Rpn13(1–128)-HTP-FLAG. Following the treatment, compound containing media was removed and cells were washed with DPBS to remove residual compound. Cells were lysed immediately using M-PER (Thermo 78501) + 1X Halt Protease Inhibitor (Thermo 1861279) with chloroalkane tagged biotin (20 μM) to alkylate any unoccupied Rpn13(1–128)-HTP-FLAG.

Western blots were run according to standard procedure and probed with streptavidin IRDye680RD (Licor 925-68079to track biotinylated HaloTag protein and anti-HaloTag antibody (Promega G928A) to track total HaloTag protein. Streptavidin was added with the secondary antibody at a 1:1000 dilution in Intercept buffer supplemented with 0.2% Tween-20 and 0.1% Sodium Dodecyl Sulfate (SDS). Phosphate buffered saline with 0.2% Tween-20 and 0.1% SDS was also used as the wash buffer following streptavidin addition. Lower biotin signal indicates more complete alkylation of Rpn13(1–128)-HTP-FLAG by the test compounds.

### BRD Degradation Assay

Degradation of BRD2 and BRD4 by the Halo-UIDs was assayed by Western blot. Cells were plated in a 6 well dish in complete media lacking antibiotics so that they would be 90% confluent at the beginning of the assay. Cells were treated with WJ701-706 (10 μM) in complete media lacking antibiotics for 30 minutes, during which time the compounds alkylated Rpn13(1–128)-HTP-FLAG or HTP-FLAG. Following the 30-minute alkylation, the compound containing media was removed and cells were washed with DPBS. Fresh media was added, and the cells were incubated at 37 °C for four hours to allow for ubiquitin-independent degradation. As a positive control for BRD degradation, cells were treated with MZ1 (1 μM) for the full 4.5-hour assay; not washout step was completed. Cells were then collected, lysed, and analyzed by Western blot. Blots were probed with anti-BRD2 antibody (CST 5848) or anti-BRD4 antibody (CST 13440) and anti-vinculin antibody (Sigma SAB400080), as a loading control.

### TMT quantitative proteomics and mass spectrometry

Protein samples in 100 uL of M-PER (Thermo 78501) + 1X Halt Protease Inhibitor (Thermo 1861279), were precipitated overnight at −20 °C using 4X ice cold acetone (v/v), washed 4 times with ice cold acetone and then allowed to air dry. The protein pellets were subsequently re-solubilized in 40 uL of 5% SDS (v/v) and assayed for protein using the PierceTM BCA protein assay kit (Thermo Fisher Scientific, Waltham, MA). 130-170ug of protein/sample (in a total volume of 33 uL), were processed for digestion using micro-S-TrapsTM (Protifi, Huntington, NY) according to manufacturer instructions. Briefly, proteins in 5% SDS were reduced with 1.43μL of 120mM Tris (2-carboxyethyl) phosphine hydrochloride (TCEP) at 56 °C for 20 minutes, followed by alkylation using 1.43μL of 500 mM methyl methanethiosulfonate (MMTS) for 20 minutes at ambient temperature. Finally, 8 μg of sequencing-grade trypsin (Promega) was added and the mixture was incubated for 1 hour at 47 °C. Following this incubation, 40 μL of 50 mM triethylammonium bicarbonate (TEAB) was added to the S-TrapTM and the peptides were eluted using centrifugation. Elution was repeated once. A third elution using 35μL of 50% acetonitrile (ACN) was also performed and the eluted peptides dried under a vacuum. The peptides were then resolubilized in 30 μL of 50 mM TEAB pH 8.5 and assayed using the PierceTM Quantitative Fluorometric Peptide Assay (Thermo Fisher Scientific, Waltham, MA). Fifty micrograms of peptides/sample were labeled with TMT labels (16-plex) according to the manufacturer’s instructions (Thermo Fisher Scientific, Waltham, MA), and pooled. The pooled, plexed samples were then dried under vacuum, resolubilized in 400 uL of 1% TFA (pH <3), desalted using Waters Oassis® OASIS HLB 1cc solid phase extraction cartridges (Waters, Milldford, MA), and then dried under vacuum one last time. The dried labelled peptide plexed sample was then resuspended in 400uL of 1 M TEAB and fractionated using an Agilent 1100 HPLC system and high pH reversed phase chromatography on a Zorbax Eclipse XDB-C18 column (4.6 x 150mm, 5-micron) from Agilent (Santa Clara, CA). Sixty fractions were collected at 0.5 mL/min over 105 min using a gradient of 0-5% solvent B in 10 min, 5-35% solvent B in 60 min, 35-70% solvent B in 15 min, a 10 min hold of 70% solvent B, and finally a return to 5% solvent B in 10 min. Solvent A consisted of 100mM TEAB and solvent B consisted of ACN. The 60 fractions were dried vacuum, concatenated into 5 fractions, and each peptide fraction was then cleaned up using 2 ug capacity C18 ZipTips (Millipore, Billerica, MA) according to the manufacturer’s instructions. Dried TMT-labelled peptides were reconstituted in 5 uL of 0.1% formic acid and on-line eluted into a Fusion Tribrid mass spectrometer (Thermo Scientific, San Jose, CA) from an EASY PepMapTM RSLC C18 column (2μm, 100Å, 75μm x 50cm, Thermo Scientific, San Jose, CA), using a gradient of 5-25% solvent B (80/20 acetonitrile/water, 0.1% formic acid) in 180 minutes, followed by 25-44% solvent B in 60 minutes, 44-80% solvent B in 0.1 minutes, a 5 minute hold of 80% solvent B, a return to 5% solvent B in 0.1 minutes, and finally a 10 minute hold of solvent B. All flow rates were 250nL/minute delivered using an Easy-nLC 1000 nano liquid chromatography system (Thermo Scientific, San Jose, CA). Solvent A consisted of water and 0.1% formic acid. Ions were created at 2.0kV using an EASY Spray source (Thermo Scientific, San Jose, CA) held at 50°C.

A synchronous precursor selection (SPS)-MS3 mass spectrometry method was selected based on the work of Ting et al.. (47) Scans were conducted between 380-2000 m/z at a resolution of 120,000 for MS1 in the Orbitrap mass analyzer at an AGC target of 4E5 and a maximum injection of 50 msec. We then performed CID in the linear ion trap of peptide monoisotopic ions with charge 2-8 above an intensity threshold of 5E3, using a quadrupole isolation of 0.7 m/z and a CID energy of 35%. The ion trap AGC target was set to 1.0E4 with a maximum injection time of 50 msec. Dynamic exclusion duration was set at 60 seconds and ions were excluded after one time within the +/− 10ppm mass tolerance window. The top 10 MS2 ions in the ion trap between 400-1200 m/z were then chosen for HCD at 65% energy. Detection occurred in the Orbitrap at a resolution of 60,000 and an AGC target of 1E5 and an injection time of 120msec (MS3). All scan events occurred within a 3-second specified cycle time.

### Proteomic Data Processing and Statistical Analysis

Quantitative analysis of the TMT pro 16plex experiment was performed simultaneously to protein identification using Proteome Discoverer 2.5 (PD) software. The precursor and fragment ion mass tolerances were set to 10 ppm, 0.6 Da, respectively), enzyme was Trypsin with a maximum of 2 missed cleavages and FASTA files for Uniprot Human (UP000005640, downloaded on 9/14/2020) proteome, the sequence of Rpn13(1–128)-HTP-FLAG and common contaminants were used in SEQUEST searches. The impurity correction factors obtained from Thermo Fisher Scientific for the TMT 16plex kit was included in the search and quantification. The following settings were used to search the data; dynamic modifications; Oxidation / +15.995Da (M), Deamidated / +0.984 Da (N, Q), N-Terminal modification of Acetyl / +42.011 Da (N-Terminus), Met-loss / −131.040 Da (M), Met-loss+Acetyl / −89.030 Da (M) and static modifications of TMTpro / +304.207 Da (Any N-Terminus, K) and MMTS +45.988 Da (C). Scaffold (version Scaffold_5.0.1, Proteome Software Inc., Portland, OR) was used to validate MS/MS based peptide and protein identifications. Peptide identifications were accepted if they had FDR < 1.0% by the Percolator posterior error probability calculation. (48) Protein identifications were accepted if they achieved an FDR < 1.0% and contained at least 2 identified peptides. Protein probabilities were assigned by the Protein Prophet algorithm.(49) Proteins sharing significant peptide evidence were grouped into clusters. 6175 proteins were clustered into 3753 proteins groups. ANOVA was used to identify proteins differentially regulated across all treatment conditions. Benjamini-Hochberg method was used to apply FDR<0.01. The proteins that passed these criteria were further analyzed by pairwise comparisons of select treatment conditions with t-test. All proteomics data analysis was performed at Bioinformatics and Statistics Core (Facility RRID:SCR_023048).

### BRD Recruitment Assay

Recruitment of BRD2 and BRD4 to Rpn13(1–128)-HTP-FLAG upon Halo-UID treatment was assayed by co-immunoprecipitation. Rpn13(1–128)-HTP-FLAG was precipitated using M2 anti-FLAG resin (Sigma #M8823) and the precipitate was blotted for BRD2 and BRD4.

Rpn13(1–128)-HTP-FLAG cells were pretreated with bortezomib (10 uM) (Tocris 7282) for 1 hour followed by Halo-UID (10 uM) treatment for 30 minutes. Compounds were washed out and the cells were incubated for 2 hours to allow complexes to form. Cells were then collected and lysed in M-PER (Thermo 78501) + 1X Halt Protease Inhibitor (Thermo 1861279) according to the manufacturer’s protocol. Lysates were clarified by centrifugation at 14,000 RCF, 15 min, 4°C. 50 uL of packed M2 resin was equilibrated into TBS. 100 uL of concentrated lysate was then added and tumbled with the resin for 30 minutes at 4°C. The resin was washed 3x with 500 uL of TBS, and the bound proteins were eluted by heating the resin in 1x Laemmli buffer at 95°C for 10 minutes. Samples of the naïve lysate were also prepared for Western blot analysis. Membranes were blotted for the FLAG epitope tag (BioLegend 637301) and either BRD2 (CST 5848) or BRD4 (CST 13440). Co-precipitation of BRD2 and BRD4 with Rpn13(1–128)-HTP-FLAG after Halo-UID treatment indicated the formation of a ternary complex.

### Proteasome Dependence Assay

The proteasome-dependence of WJ704-706 degradation of BRD2 was assayed by co-treatment with bortezomib (Tocris 7282), a proteasome inhibitor. Cells were plated in a 6 well dish in complete media lacking antibiotics so that they would be 90% confluent at the beginning of the assay. Cells were pre-treated with either bortezomib (10 μM) or DMSO for 1 hour. Following the pre-treatment, the cells were treated with WJ704-706 (10 μM) with or without bortezomib for 30 minutes to alkylate the HaloTag protein. The media was then removed, and the cells were washed with DPBS. The media was replaced with complete media with either bortezomib or DMSO. After four hours, the cells were collected, lysed, and analyzed by Western blot. Blots were probed with anti-BRD2 antibody (CST 5848) and anti-vinculin antibody (Sigma SAB400080), as a loading control.

### Ubiquitin Independence Assay

The ubiquitin-dependence of WJ704-706 degradation of BRD2 was assayed by co-treatment with TAK-243 (MCE HY-100487), also known as MLN7243, a ubiquitin activating enzyme inhibitor. Cells were plated in a 6 well dish in complete media lacking antibiotics so that they would be 90% confluent at the beginning of the assay. Cells were pre-treated with either TAK-243 (10 μM) or DMSO for 30 minutes. Following the pre-treatment, the cells were treated with WJ704-706 (10 μM) with or without TAK-243 for 30 minutes to alkylate the HaloTag protein. The media was then removed, and the cells were washed with DPBS. The media was replaced with complete media with either TAK-243 or DMSO. After four hours, the cells were collected, lysed, and analyzed by western blot. Blots were probed with anti-BRD2 antibody (CST 5848) and anti-vinculin antibody (Sigma SAB400080), as a loading control.

### NAE Independence Assay

The neddylation-dependence of WJ704-706 degradation of BRD2 was assayed by co-treatment with TAK-924 (Tocris 6499), also known as MLN4924, a nedd activating enzyme inhibitor. Cells were plated in a 6 well dish in complete media lacking antibiotics so that they would be 90% confluent at the beginning of the assay. Cells were pre-treated with either TAK-924 (10 µM) or DMSO for 1 hour. Following the pre-treatment, the cells were treated with WJ704-706 (10 µM) with or without TAK-924 for 30 minutes to alkylate the HaloTag protein. The media was then removed, and the cells were washed with DPBS. The media was replaced with complete media with either 10 TAK-924 or DMSO. After four hours, the cells were collected, lysed, and analyzed by western blot. Blots were probed with anti-BRD2 antibody (CST 5848) and anti-vinculin antibody (Sigma SAB400080), as a loading control.

### JQ1 Competition Assay

The specificity of BRD2 recruitment to the proteasome was assayed by co-treatment with JQ1 (Tocris 4499), the BET inhibitor that was used as a TP ligand in the Halo-UIDs. Cells were plated in a 6 well dish in complete media lacking antibiotics so that they would be 90% confluent at the beginning of the assay. Cells were pre-treated with either JQ1 (10 µM) or DMSO for 1 hour. Following the pre-treatment, the cells were treated with WJ704-706 (10 µM) with or without JQ1 for 30 minutes to alkylate the HaloTag protein. The media was then removed, and the cells were washed with DPBS. The media was replaced with complete media with either JQ1 or DMSO. After four hours, the cells were collected, lysed, and analyzed by Western blot. Blots were probed with anti-BRD2 antibody (CST 5848) and anti-vinculin antibody (Sigma SAB400080), as a loading control.

## Data Availability Statement

The data to support the conclusions in this study are presented in the main text as well as the supplementary information.

## Acknowledgements

We thank Drs. Gogce Crynen and George Tsaprailis (UF Scripps Proteomics Core) for help with carrying out and interpreting the proteomics experiments.

## Author Contributions

T.K. and M.B, conceptualization; M.B., W.G., B.-B.S. I.J. and J.T. investigation and data curation; T.K and M.B. writing, T.K. funding acquisition.

