## Supplementary material for "Chemically Induced Degradation of Native Proteins by Direct Recruitment to the 26S Proteasome": SI

### Contents:

**General Synthesis Methods……………………………………………………………………………2**

**Chemical Characterization of Halo-UIDs**

WJ701……………………………………………………………………………………...4
WJ702……………………………………………………………………………………...4
WJ703……………………………………………………………………………………...4
WJ704
WJ705……………………………………………………………………………………...5
WJ706……………………………………………………………………………………...5

**NMR Spectra**

WJ701……………………………………………………………………………………...6
WJ702……………………………………………………………………………………...7
WJ703……………………………………………………………………………………...8
WJ704……………………………………………………………………………………...9
WJ705……………………………………………………………………………………10
WJ706……………………………………………………………………………………11

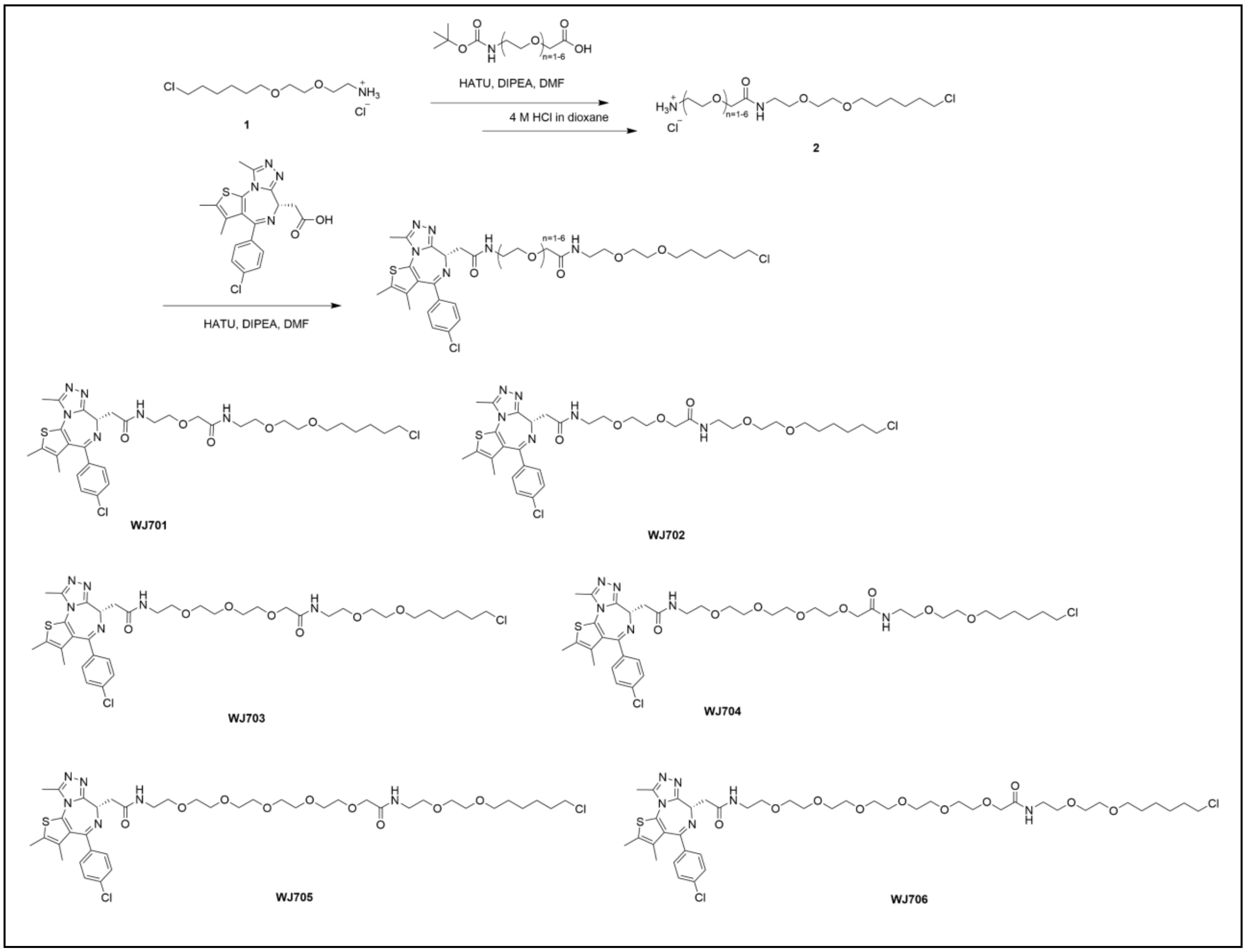


**General information.**

Chemical reagents were purchased from Fisher Scientific, Millipore Sigma, Alfa and Acros. $^{1}H$ and $^{13}C$ NMR spectra were obtained with Bruker AV600 NMR Spectrometer with a CryoProbe. Chemical shifts are reported in δ (ppm) units and $^{13}C$ and $^{1}H$ signals from deuterated solvents were used as references. 60 F254 (Merck) silica plate was used in thin layer chromatography (TLC) analysis.

Agilent 1100 series HPLC equipped with Agilent ZORBAX SB-C18 column (Part number: 861953-902, 4.6*100 mm. 3.5 µm) and an electrospray ionization (ESI) source (Agilent Technologies 6120 Quadrupole) was used to obtain mass spectra for small molecules. The following methods were used to analyze the purity of the samples in the LC-MS instrument. Solvent A contains 100% water (0.1% formic acid) and solvent B contains 100% acetonitrile (0.1% formic acid).

| Time (min) | A [%] | B [%] | Flow [mL/Min] |
|---|---|---|---|
| 0.0 | 100.0 | 0.0 | 1.0 |
| 2.0 | 100.0 | 0.0 | 1.0 |
| 12.0 | 0.0 | 100.0 | 1.0 |
| 15.0 | 0.0 | 100.0 | 1.0 |

Small molecule column chromatography was conducted on silica gel (230-400 mesh) or a SunFire Prep C18 column (100Å, 5 µm, 19 mm X 250 mm, Waters) was used at flow rate of 9 mL/min. The UV absorbance at 214, 254, 298 nm were used to monitor the purification process. Water (with 0.1% Trifluoroacetic acid) and acetonitrile (with 0.1% Trifluoroacetic acid) were used as solvent A and B respectively. The following methods were used to purify the samples.

| Time (min) | A [%] | B [%] | Flow [mL/Min] |
|---|---|---|---|
| 0.0 | 95.0 | 5.0 | 9.0 |
| 5.0 | 95.0 | 5.0 | 9.0 |
| 60.0 | 15.0 | 85.0 | 9.0 |
| 70.0 | 5.0 | 95.0 | 9.0 |
| 80.0 | 5.0 | 95.0 | 9.0 |
| 81.0 | 95.0 | 5.0 | 9.0 |
| 90.0 | 95.0 | 5.0 | 9.0 |

The compound **1** was synthesized according to a previous publication.[1]

**The general procedure to remove the Boc protection group.**

Boc protected compound (0.5 mmol) was dissolved in 7.5 mL DCM. Then, 4 M HCL in dioxane (10 mL, 40 mmol) was added to the solution and stirred at room temperature for 2 h. After completion, the solvent was evaporated, and the crude product was precipitate with dry $Et_2O$, washed with $Et_2O$ for three times and used for next step coupling reaction.

**The general procedure for amide coupling.**

Method 1: Carboxylic acid (0.15 mmol) was dissolved in 1 mL DMF, then HATU (72 mg, 0.2 mol), N,N-Diisopropylethylamine (87 µL, 0.5 mmol) were added and stirred at room temperature for 10 min. Then, amine (0.2 mmol) was added to the solution and stirred at room temperature for 12 h. The reaction was monitored by LC/MS. After completion, the product was purified by HPLC.

**WJ701: (S)-N-(18-chloro-5-oxo-3,9,12-trioxa-6-azaoctadecyl)-2-(4-(4-chlorophenyl)-2,3,9-trimethyl-6H-thieno[3,2-f][1,2,4]triazolo[4,3-a][1,4]diazepin-6-yl)acetamide.** $^1$H NMR (600 MHz, DMSO-$d_6$) δ 8.25 (t, $J$ = 5.7 Hz, 1H), 7.68 (t, $J$ = 5.9 Hz, 1H), 7.39 – 7.37 (m, 2H), 7.34 – 7.30 (m, 2H), 4.42 (dd, $J$ = 8.3, 6.0 Hz, 1H), 3.79 (s, 2H), 3.49 (t, $J$ = 6.6 Hz, 2H), 3.43 – 3.36 (m, 4H), 3.36 – 3.31 (m, 4H), 3.31 – 3.08 (m, 8H), 2.50 (s, 3H), 2.31 (d, $J$ = 0.9 Hz, 3H), 1.57 (dq, $J$ = 8.7, 6.7 Hz, 2H), 1.51 (d, $J$ = 0.9 Hz, 3H), 1.41 – 1.32 (m, 2H), 1.30 – 1.22 (m, 2H), 1.18 (dddd, $J$ = 8.9, 7.1, 4.7, 1.5 Hz, 2H). $^{13}$C NMR (151 MHz, DMSO) δ 170.11, 169.63, 163.65, 155.52, 150.45, 137.11, 135.77, 132.69, 131.36, 130.68, 130.36, 130.08, 128.96, 70.65, 70.35, 70.32, 70.02, 69.87, 69.34, 54.23, 45.83, 38.90, 38.51, 37.96, 32.47, 29.50, 26.57, 25.37, 14.53, 13.16, 11.76. MS (ESI, positive) m/z calculated for $C_{33}H_{45}Cl_2N_6O_5S$ $[M+H]^+$ : 707.3, found: 707.3.

**WJ702: (S)-N-(21-chloro-8-oxo-3,6,12,15-tetraoxa-9-azahenicosyl)-2-(4-(4-chlorophenyl)-2,3,9-trimethyl-6H-thieno[3,2-f][1,2,4]triazolo[4,3-a][1,4]diazepin-6-yl)acetamide.** $^1$H NMR (600 MHz, DMSO-$d_6$) δ 8.29 (t, $J$ = 5.7 Hz, 1H), 7.65 (t, $J$ = 5.9 Hz, 1H), 7.49 (d, $J$ = 8.6 Hz, 2H), 7.42 (d, $J$ = 8.3 Hz, 2H), 4.52 (dd, $J$ = 8.0, 6.1 Hz, 1H), 3.88 (s, 2H), 3.63 – 3.56 (m, 6H), 3.51 – 3.41 (m, 8H), 3.37 – 3.20 (m, 8H), 2.60 (s, 3H), 2.41 (s, 3H), 1.73 – 1.65 (m, 2H), 1.62 (s, 3H), 1.52 – 1.41 (m, 2H), 1.41 – 1.33 (m, 2H), 1.33 – 1.24 (m, 2H). $^{13}$C NMR (151 MHz, DMSO) δ 170.11, 163.63, 155.53, 150.44, 137.13, 135.78, 132.69, 131.35, 130.67, 130.36, 130.08, 128.96, 70.67, 70.65, 70.41, 70.00, 69.89, 69.86, 69.68, 69.36, 54.23, 45.83, 39.08, 38.45, 37.87, 32.47, 29.51, 26.58, 25.38, 14.52, 13.16, 11.76. MS (ESI, positive) m/z calculated for $C_{35}H_{49}Cl_2N_6O_6S$ $[M+H]^+$ : 751.3, found: 751.3.

**WJ703: (S)-N-(24-chloro-11-oxo-3,6,9,15,18-pentaoxa-12-azatetracosyl)-2-(4-(4-chlorophenyl)-2,3,9-trimethyl-6H-thieno[3,2-f][1,2,4]triazolo[4,3-a][1,4]diazepin-6-yl)acetamide.** $^1$H NMR (600 MHz, DMSO-$d_6$) δ 8.29 (t, $J$ = 5.7 Hz, 1H), 7.64 (t, $J$ = 5.9 Hz, 1H), 7.51 – 7.47 (m, 2H), 7.45 – 7.40 (m, 2H), 4.52 (dd, $J$ = 8.1, 6.1 Hz, 1H), 3.88 (s, 2H), 3.63 – 3.54 (m, 10H), 3.50 – 3.40 (m, 8H), 3.35 (t, $J$ = 6.6 Hz, 2H), 3.34 – 3.18 (m, 6H), 2.60 (s, 3H), 2.41 (d, $J$ = 1.0 Hz, 3H), 1.72 – 1.65 (m, 2H), 1.62 (d, $J$ = 0.9 Hz, 3H), 1.51 – 1.44 (m, 2H), 1.36 (tdd, $J$ = 9.1, 4.4, 3.2 Hz, 2H), 1.29 (dddd, $J$ = 8.9, 7.0, 4.7, 1.5 Hz, 2H). $^{13}$C NMR (151 MHz, DMSO) δ 170.09, 169.72, 163.62, 155.54, 150.44, 137.12, 135.78, 132.69, 131.35, 130.68, 130.36, 130.08, 128.95, 70.68, 70.66, 70.39, 70.23, 70.09, 70.05, 70.01, 70.00, 69.90, 69.68, 69.35, 54.22, 45.83, 39.08, 38.45, 37.86, 32.47, 29.52, 26.58, 25.39, 14.53, 13.16, 11.76. MS (ESI, positive) m/z calculated for $C_{37}H_{53}Cl_2N_6O_7S$ $[M+H]^+$ : 795.3, found: 795.3.

**WJ704: (S)-N-(2-(2-((6-chlorohexyl)oxy)ethoxy)ethyl)-14-(2-(4-(4-chlorophenyl)-2,3,9-trimethyl-6H-thieno[3,2-f][1,2,4]triazolo[4,3-a][1,4]diazepin-6-yl)acetamido)-3,6,9,12-tetraoxatetradecanamide.** $^{1}$H NMR (600 MHz, DMSO-$d_6$) δ 8.29 (t, *J* = 5.7 Hz, 1H), 7.64 (t, *J* = 6.0 Hz, 1H), 7.50 – 7.47 (m, 2H), 7.44 – 7.40 (m, 2H), 4.52 (dd, *J* = 8.2, 6.0 Hz, 1H), 3.87 (s, 2H), 3.61 (t, *J* = 6.6 Hz, 2H), 3.58 – 3.52 (m, 12H), 3.51 – 3.48 (m, 2H), 3.47 – 3.41 (m, 6H), 3.36 (t, *J* = 6.5 Hz, 2H), 3.33 – 3.18 (m, 6H), 2.60 (s, 3H), 2.41 (d, *J* = 1.0 Hz, 3H), 1.69 (dq, *J* = 8.7, 6.7 Hz, 2H), 1.62 (d, *J* = 0.9 Hz, 3H), 1.52 – 1.44 (m, 2H), 1.36 (tdd, *J* = 9.0, 4.3, 3.2 Hz, 2H), 1.29 (dddd, *J* = 8.9, 7.0, 4.7, 1.5 Hz, 2H). $^{13}$C NMR (151 MHz, DMSO) δ 170.08, 169.71, 163.62, 150.44, 137.12, 135.78, 132.69, 131.35, 130.68, 130.35, 130.08, 128.95, 70.69, 70.66, 70.38, 70.37, 70.27, 70.23, 70.10, 70.01, 69.90, 69.68, 69.35, 54.22, 45.83, 38.44, 37.87, 32.48, 29.52, 29.51, 26.58, 25.39, 14.52, 13.15, 11.76. MS (ESI, positive) m/z calculated for $C_{39}H_{57}Cl_2N_6O_8S$ $[M+H]^+$ : 839.3, found: 839.3.

**WJ705: (S)-N-(2-(2-((6-chlorohexyl)oxy)ethoxy)ethyl)-17-(2-(4-(4-chlorophenyl)-2,3,9-trimethyl-6H-thieno[3,2-f][1,2,4]triazolo[4,3-a][1,4]diazepin-6-yl)acetamido)-3,6,9,12,15-pentaoxaheptadecanamide.** $^{1}$H NMR (600 MHz, DMSO-$d_6$) δ 8.29 (t, *J* = 5.7 Hz, 1H), 7.66 – 7.61 (m, 1H), 7.51 – 7.47 (m, 2H), 7.45 – 7.41 (m, 2H), 4.52 (dd, *J* = 8.2, 6.0 Hz, 1H), 3.86 (s, 2H), 3.61 (t, *J* = 6.6 Hz, 2H), 3.58 – 3.48 (m, 18H), 3.47 – 3.41 (m, 6H), 3.36 (t, *J* = 6.6 Hz, 2H), 3.34 – 3.18 (m, 6H), 2.61 (s, 3H), 2.41 (d, *J* = 0.9 Hz, 3H), 1.69 (dt, *J* = 14.5, 6.7 Hz, 2H), 1.62 (d, *J* = 0.9 Hz, 3H), 1.47 (dq, *J* = 8.0, 6.7 Hz, 2H), 1.41 – 1.33 (m, 2H), 1.30 (dddd, *J* = 8.9, 7.1, 4.7, 1.5 Hz, 2H). $^{13}$C NMR (151 MHz, DMSO) δ 170.07, 169.71, 163.64, 155.53, 150.47, 137.10, 135.80, 132.68, 131.39, 130.70, 130.36, 130.09, 128.95, 70.69, 70.66, 70.37, 70.26, 70.23, 70.10, 70.01, 69.99, 69.91, 69.69, 69.35, 54.21, 45.83, 39.10, 38.43, 37.85, 32.48, 29.52, 26.58, 25.39, 14.52, 13.15, 11.76. MS (ESI, positive) m/z calculated for $C_{41}H_{61}Cl_2N_6O_9S$ $[M+H]^+$ : 883.4, found: 883.4.

**WJ706: (S)-N-(2-(2-((6-chlorohexyl)oxy)ethoxy)ethyl)-20-(2-(4-(4-chlorophenyl)-2,3,9-trimethyl-6H-thieno[3,2-f][1,2,4]triazolo[4,3-a][1,4]diazepin-6-yl)acetamido)-3,6,9,12,15,18-hexaoxaicosanamide.** $^{1}$H NMR (600 MHz, DMSO-$d_6$) δ 8.29 (t, *J* = 5.7 Hz, 1H), 7.63 (t, *J* = 5.9 Hz, 1H), 7.51 – 7.47 (m, 2H), 7.45 – 7.40 (m, 2H), 4.52 (dd, *J* = 8.2, 6.0 Hz, 1H), 3.86 (s, 2H), 3.61 (t, *J* = 6.6 Hz, 2H), 3.58 – 3.48 (m, 22H), 3.47 – 3.41 (m, 6H), 3.36 (t, *J* = 6.6 Hz, 2H), 3.32 – 3.19 (m, 6H), 2.61 (s, 3H), 2.41 (s, 3H), 1.69 (dt, *J* = 14.6, 6.7 Hz, 2H), 1.62 (s, 3H), 1.48 (dt, *J* = 14.4, 6.6 Hz, 2H), 1.37 (dddd, *J* = 9.1, 7.3, 4.5, 3.1 Hz, 2H), 1.30 (dddd, *J* = 8.9, 7.1, 4.8, 1.5 Hz, 2H). $^{13}$C NMR (151 MHz, DMSO) δ 170.07, 169.71, 163.64, 155.53, 155.52, 150.47, 137.10, 135.80, 132.67, 131.39, 130.69, 130.36, 130.10, 128.95, 70.69, 70.66, 70.37, 70.26, 70.23, 70.10, 70.01, 69.99, 69.91, 69.90, 69.69, 69.36, 54.21, 45.83, 39.10, 38.43, 37.85, 32.48, 29.52, 26.58, 25.39, 14.52, 13.15, 11.75. MS (ESI, positive) m/z calculated for $C_{43}H_{65}Cl_2N_6O_{10}S$ $[M+H]^+$ : 927.4, found: 927.4.

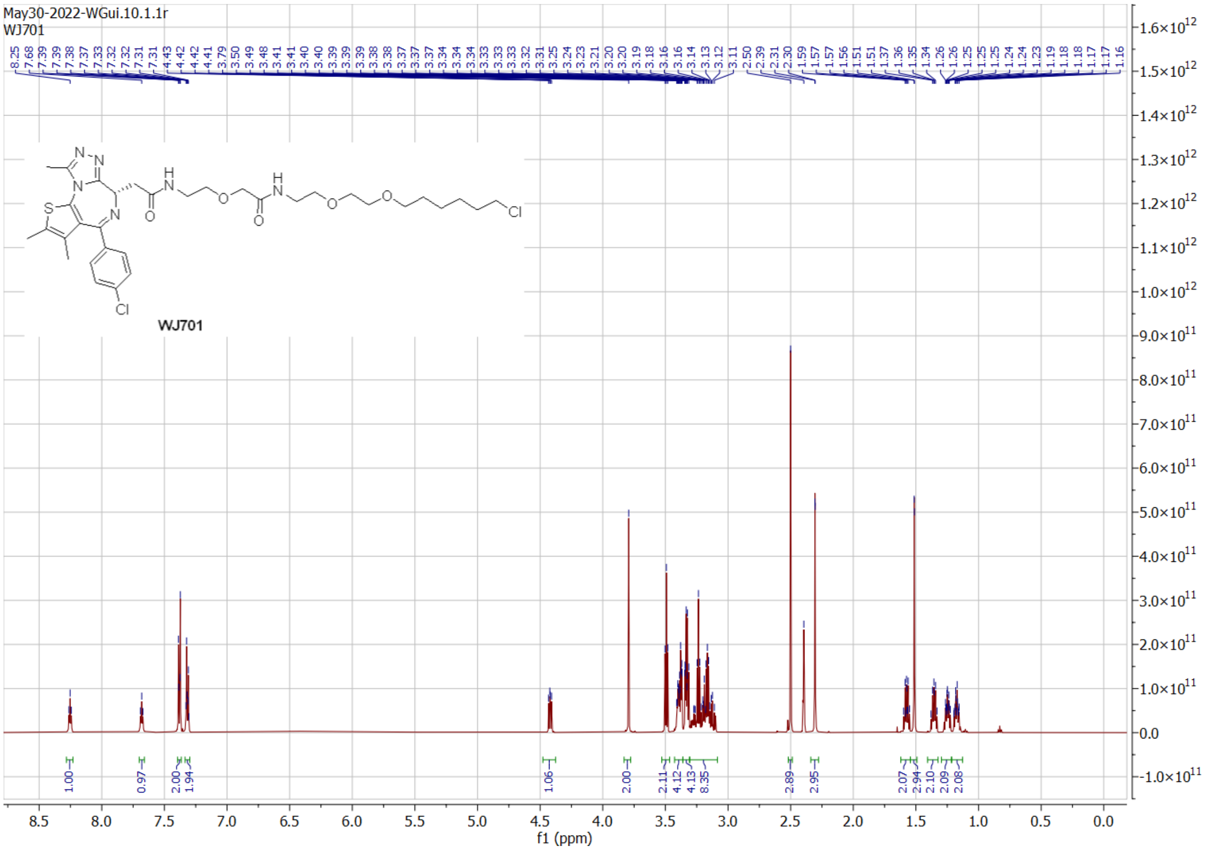
May30-2022-WGui.10.1.1r
WJ701
WJ701
f1 (ppm)

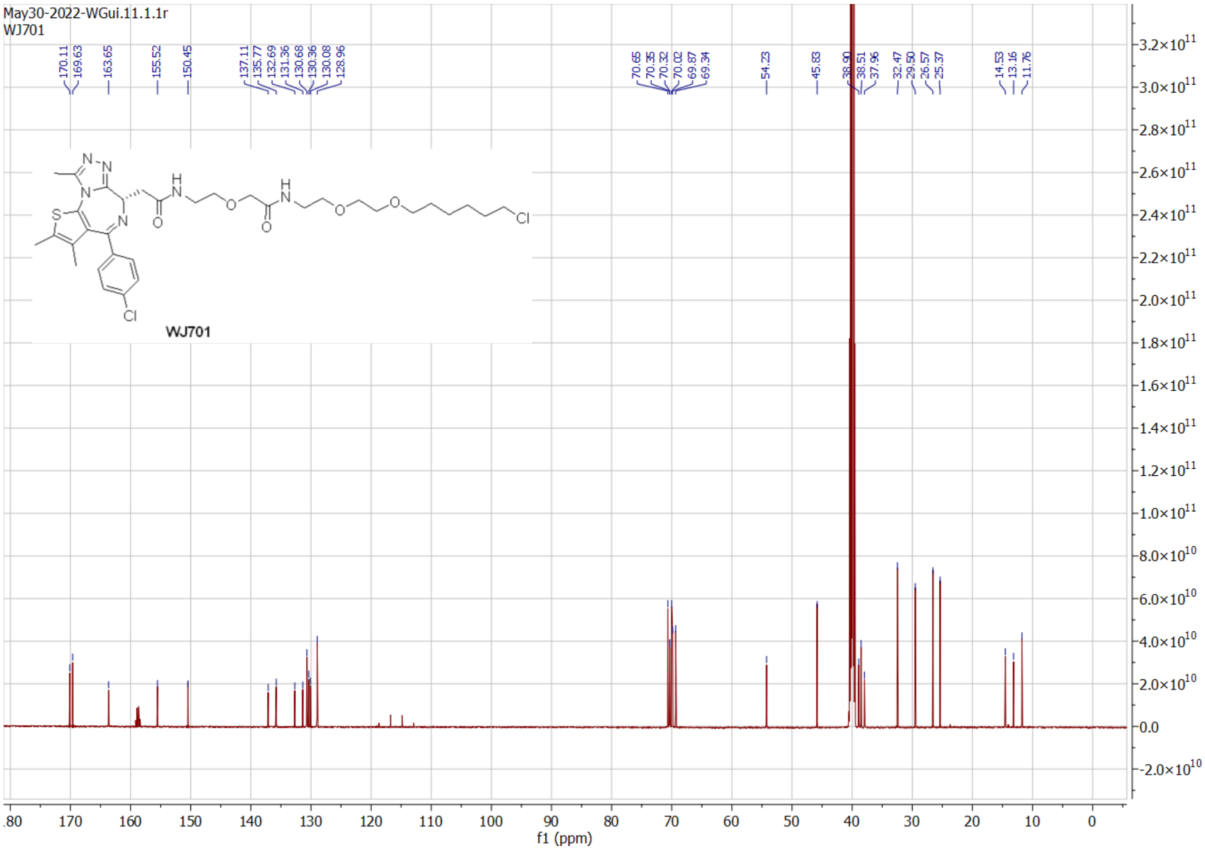
May30-2022-WGui.11.1.1r
WJ701
WJ701
f1 (ppm)

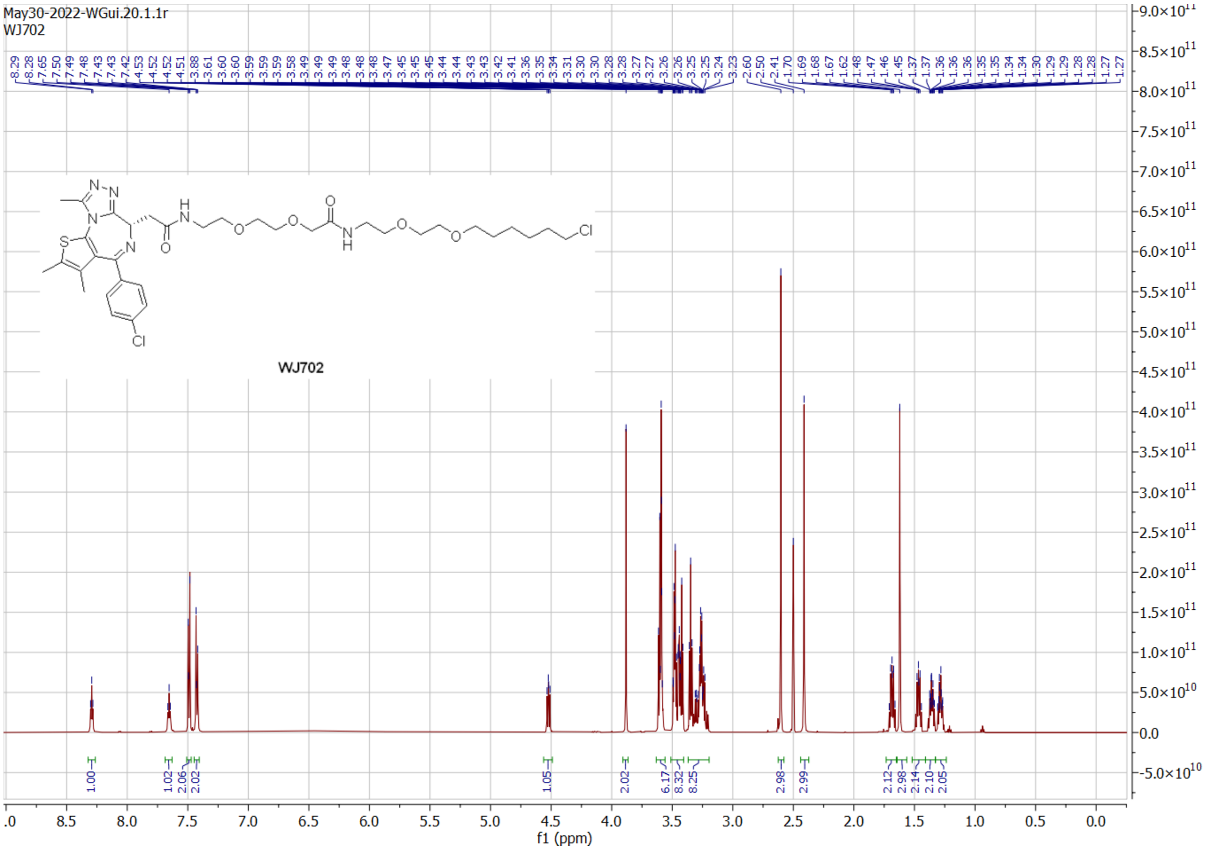
May30-2022-WGui.20.1.1r
WJ702
8.29 8.28 7.65 7.50 7.49 7.48 7.43 7.43 7.43 7.42 4.53 4.52 4.52 4.51 3.88 3.61 3.60 3.60 3.59 3.59 3.59 3.58 3.49 3.49 3.49 3.48 3.48 3.48 3.47 3.45 3.45 3.45 3.44 3.44 3.44 3.43 3.43 3.43 3.42 3.41 3.36 3.35 3.34 3.31 3.30 3.30 3.28 3.28 3.27 3.27 3.26 3.26 3.25 3.25 3.24 3.23 2.60 2.50 2.41 1.70 1.69 1.68 1.67 1.62 1.48 1.47 1.46 1.45 1.37 1.37 1.36 1.36 1.36 1.35 1.35 1.34 1.34 1.30 1.29 1.29 1.28 1.28 1.27 1.27
WJ702
1.00 1.02 2.06 2.02 1.05 2.02 6.17 8.32 8.25 2.98 2.99 2.12 2.98 2.14 2.10 2.05
f1 (ppm)

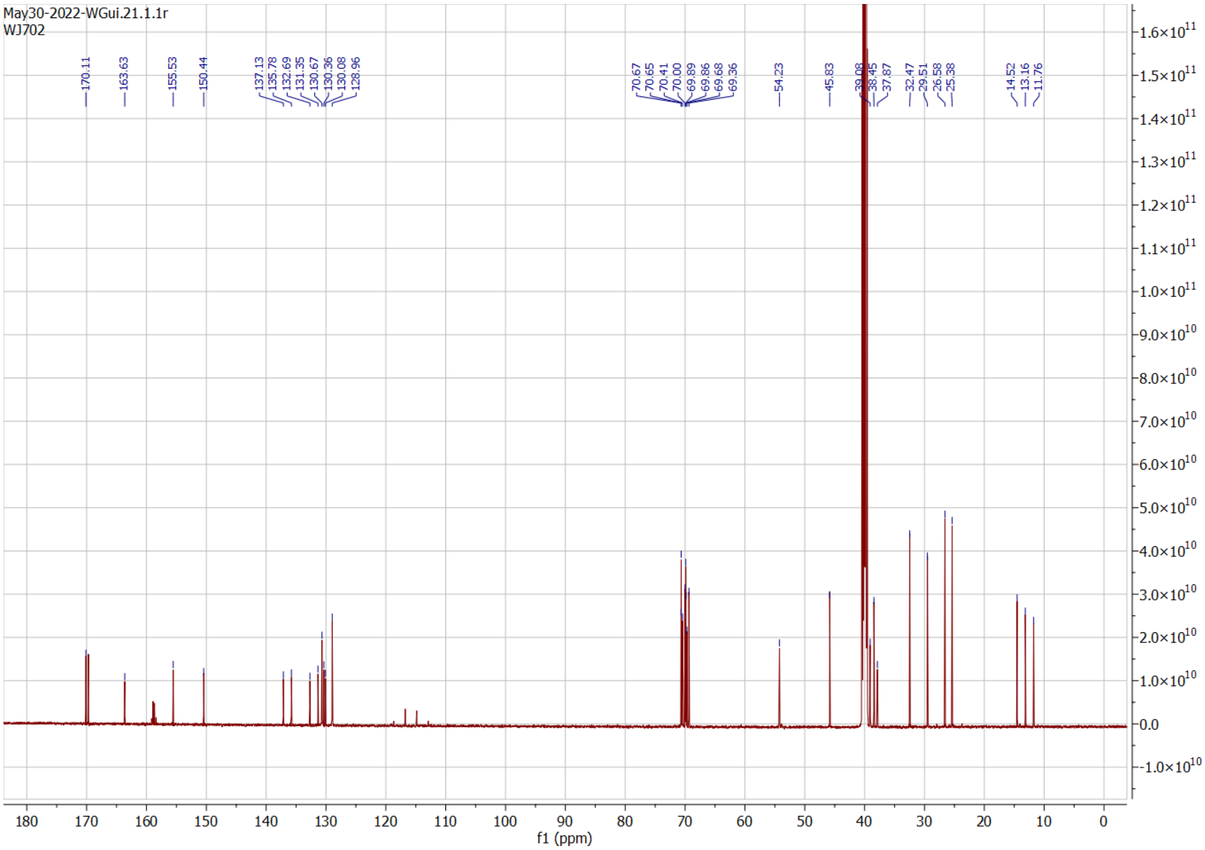
May30-2022-WGui.21.1.1r
WJ702
170.11 163.63 155.53 150.44 137.13 135.78 132.69 131.35 130.67 130.36 130.08 128.96 70.67 70.65 70.41 70.00 69.89 69.86 69.68 69.36 54.23 45.83 39.08 38.45 37.87 32.47 29.51 26.58 25.38 14.52 13.16 11.76
f1 (ppm)

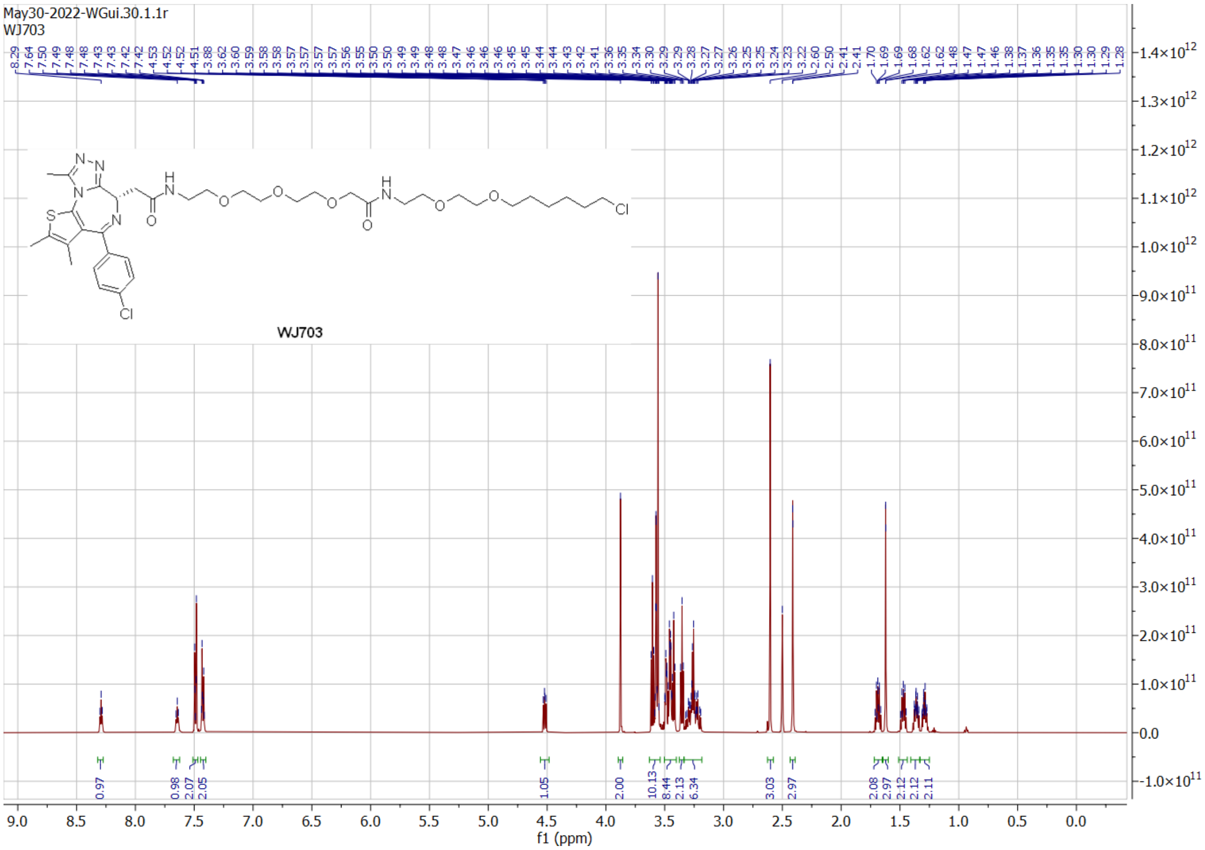
May30-2022-WGui.30.1.1r
WJ703
WJ703
f1 (ppm)

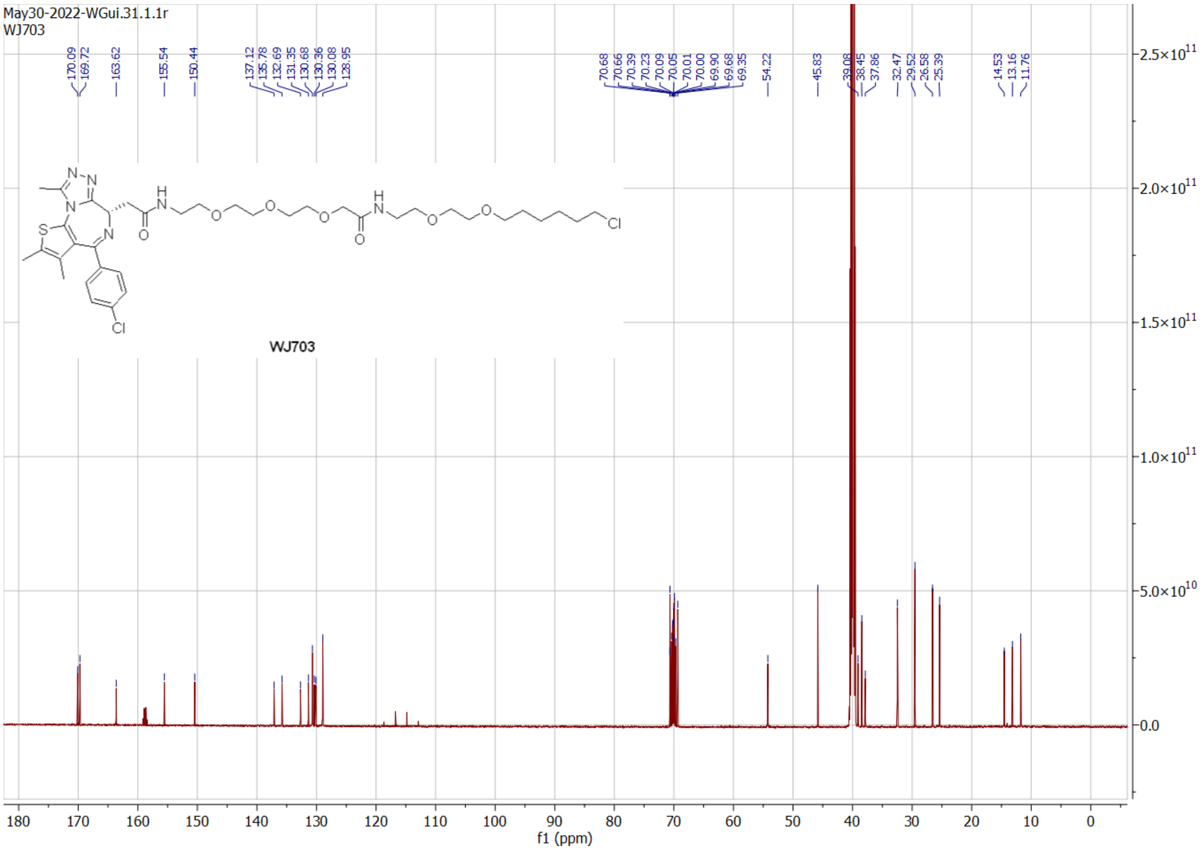
May30-2022-WGui.31.1.1r
WJ703
WJ703
f1 (ppm)

May30-2022-WGui.40.1.1r
WJ704

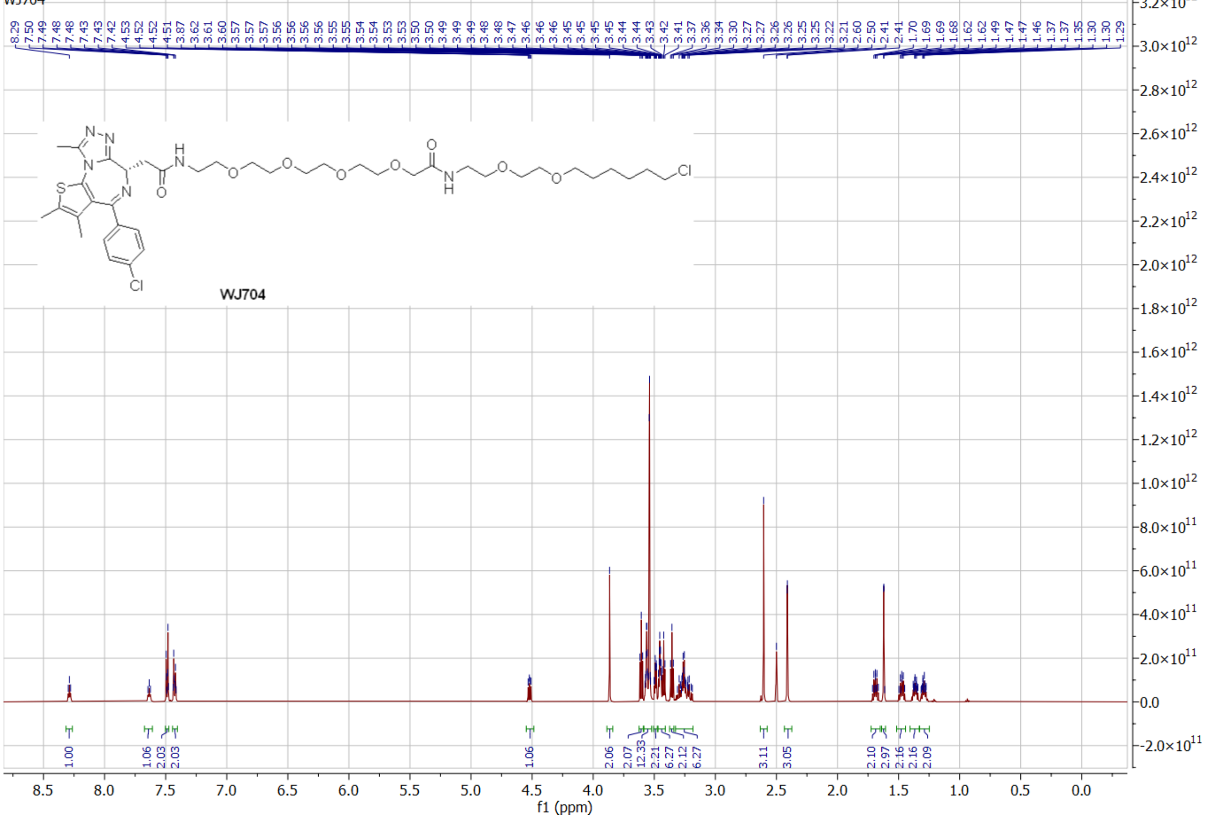


May30-2022-WGui.41.1.1r
WJ704

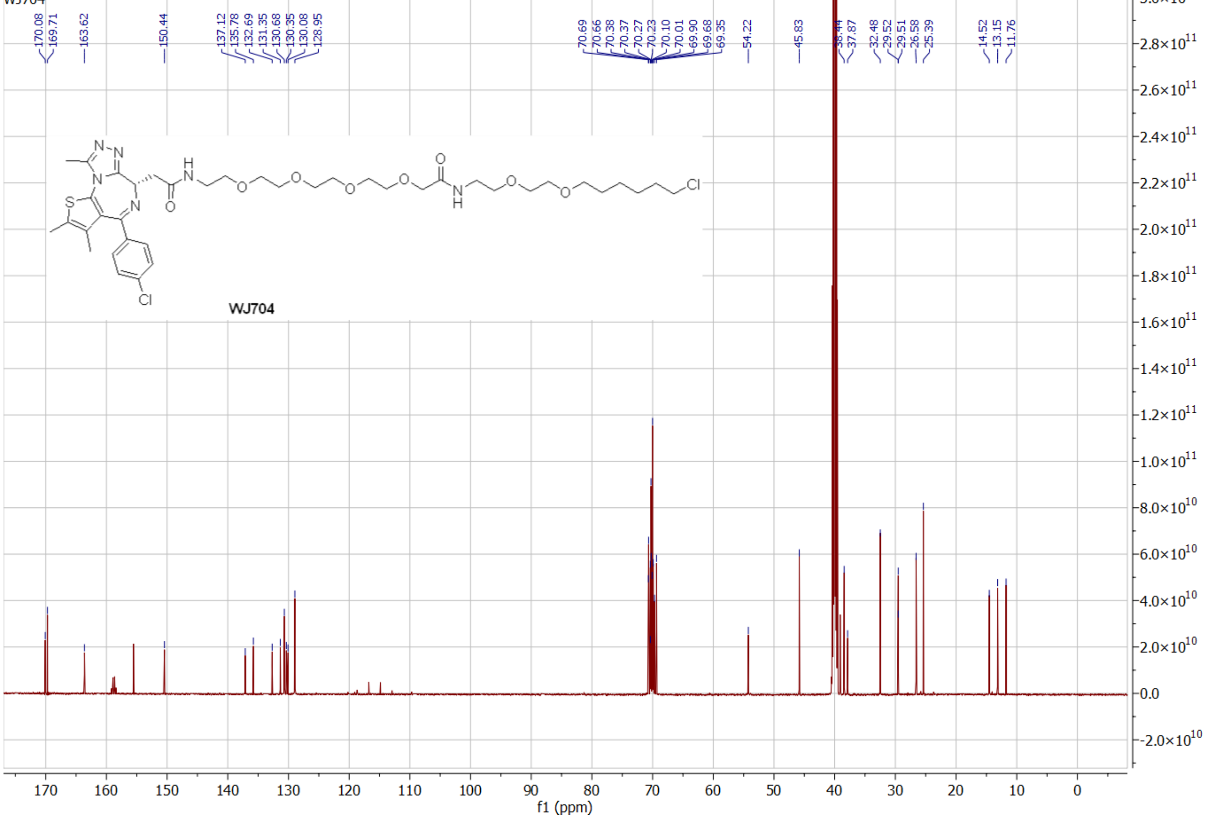

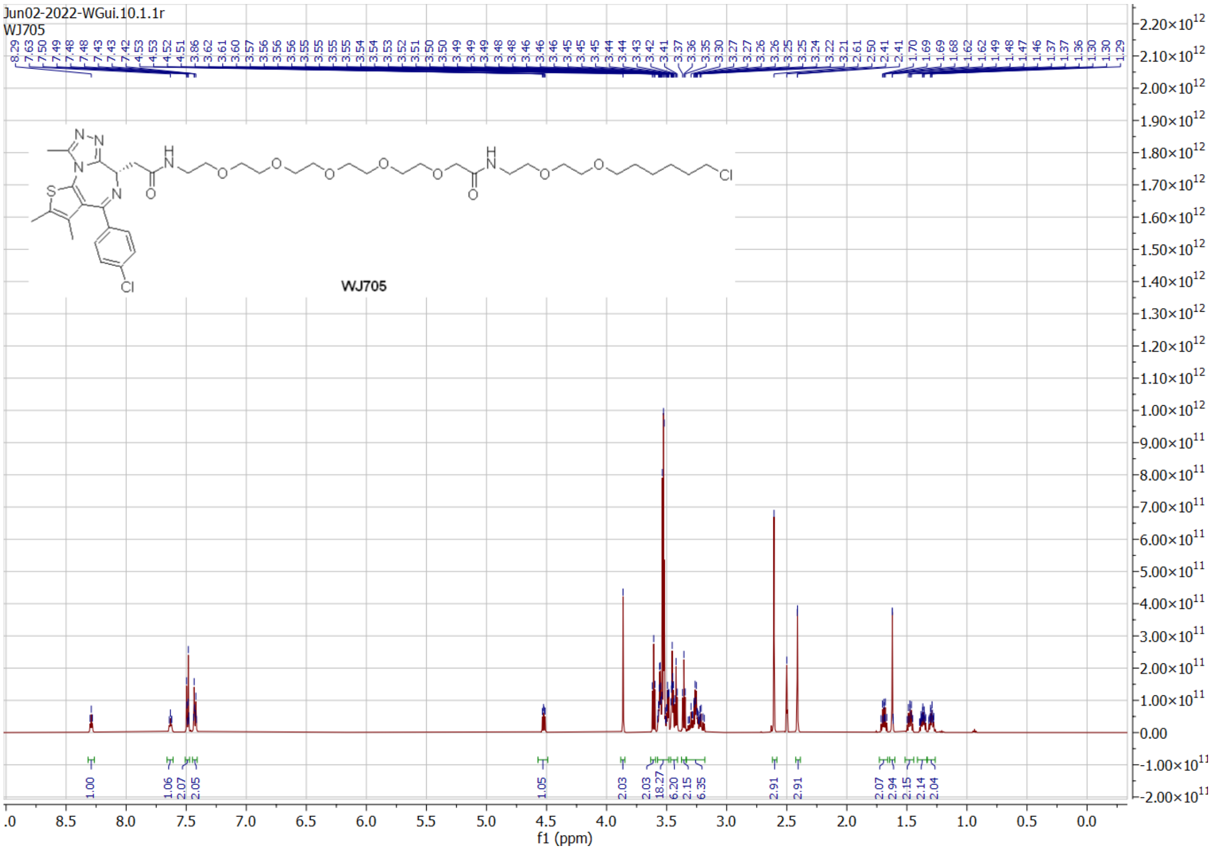
Jun02-2022-WGui.10.1.1r
WJ705
WJ705
f1 (ppm)

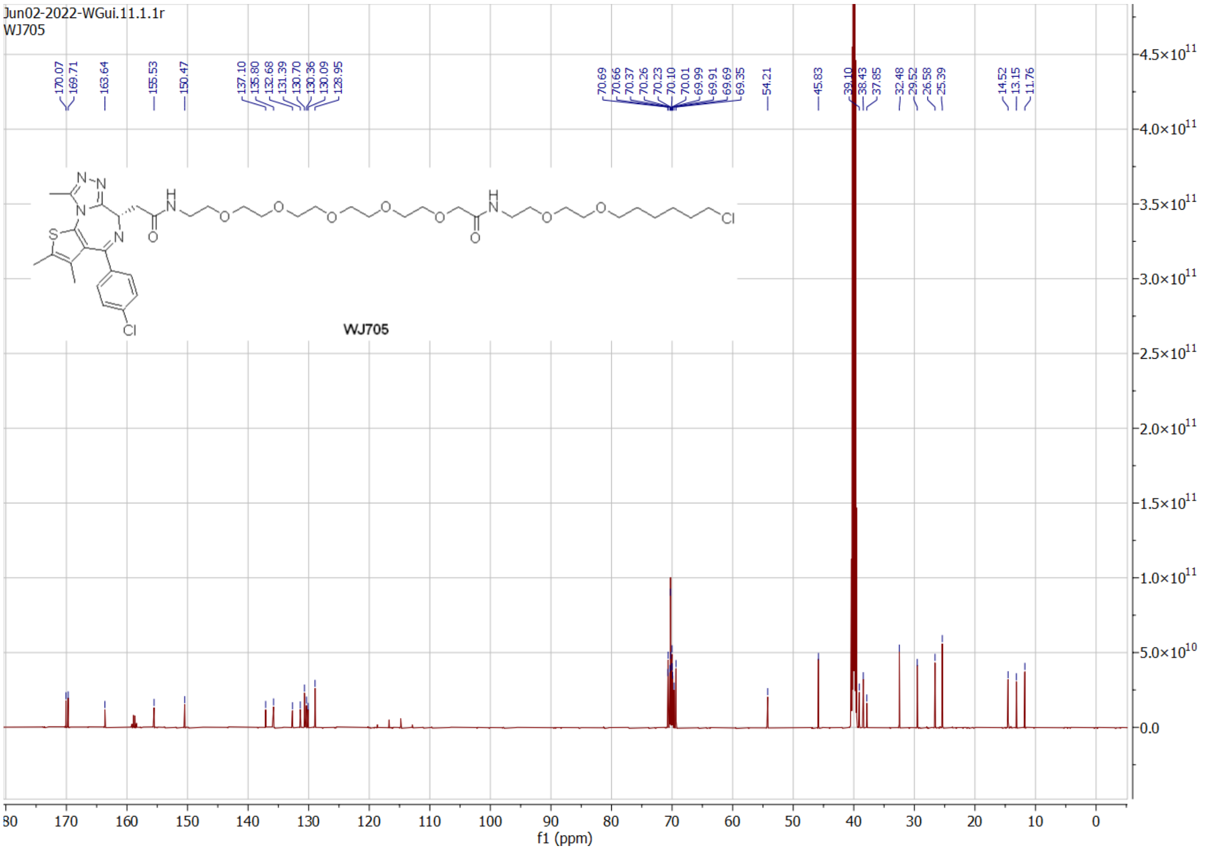
Jun02-2022-WGui.11.1.1r
WJ705
WJ705
f1 (ppm)

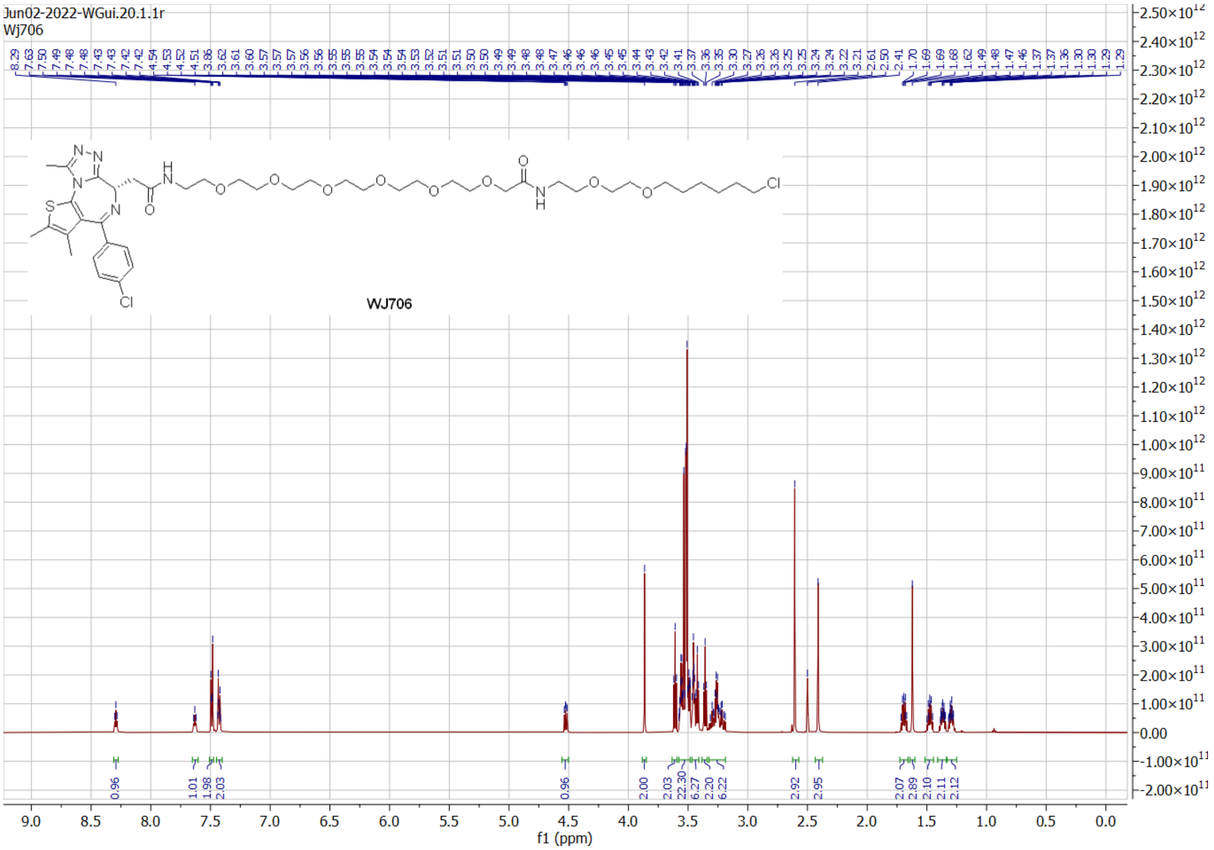

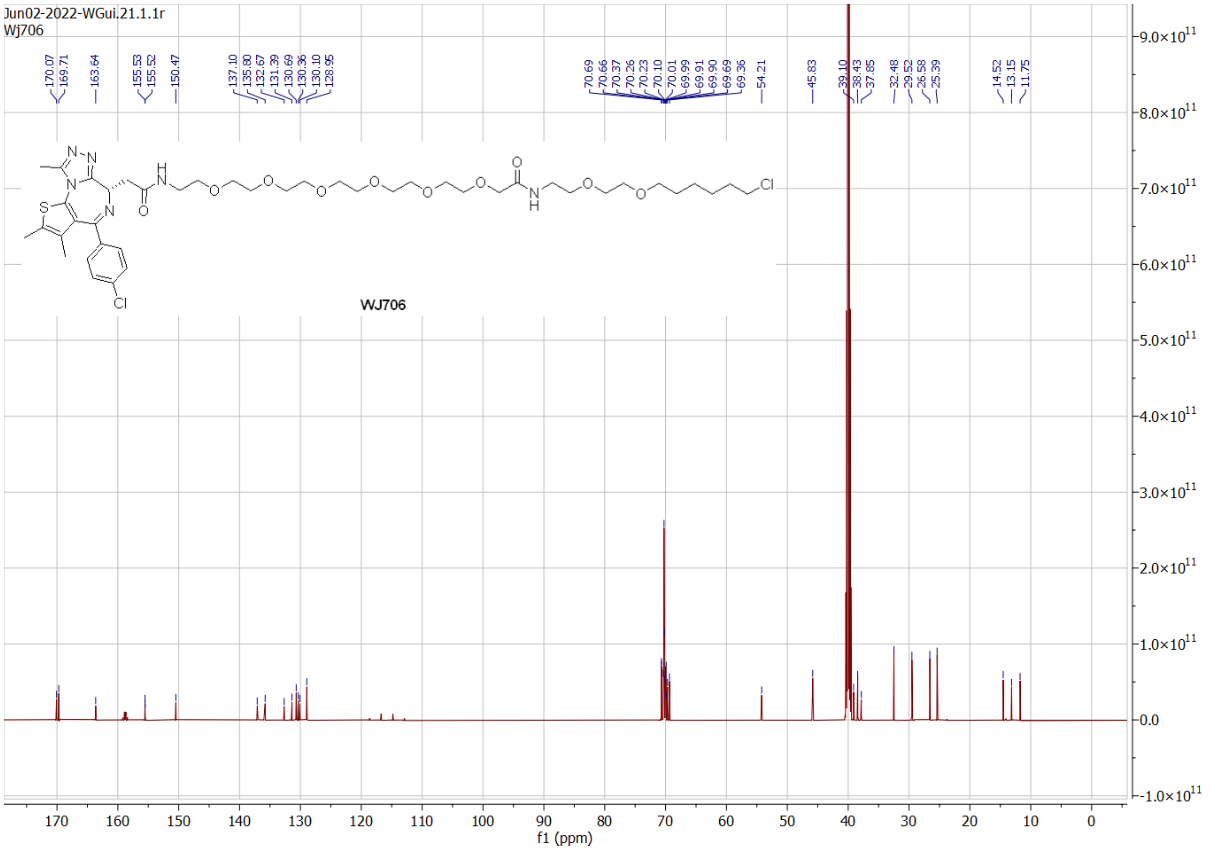

[1] L. Peraro, K. L. Deprey, M. K. Moser, Z. Zou, H. L. Ball, B. Levine, J. A. Kritzer, *J. Am. Chem. Soc.* **2018**, *140*, 11360-11369.
